# Impact of recreational fisheries management on fish biodiversity in gravel pit lakes with contrasts to unmanaged lakes

**DOI:** 10.1101/419994

**Authors:** S. Matern, M. Emmrich, T. Klefoth, C. Wolter, N. Wegener, R. Arlinghaus

## Abstract

Gravel pit lakes constitute novel ecosystems that can be colonized by fishes through natural or anthropogenic pathways. Many of these man-made lakes are used by recreational anglers and experience regular fish stocking. Recreationally unmanaged gravel pits may also be affected by fish introductions, e.g., through illegal fish releases, thereby contributing to the formation of site-specific communities. Our objective was to compare the fish biodiversity in gravel pit lakes with and without the recent influence of recreational fisheries management. We sampled 23 small (< 20 ha) gravel pit lakes (16 managed and 7 unmanaged) in north-western Germany and compared fish community and diversity metrics obtained using littoral electrofishing and multimesh gillnet catch per unit effort data. Given the size of the lakes we sampled we expected species poor communities and elevated fish diversity in the managed systems due to stocking. The two lake types were primarily mesotrophic and did not differ in key abiotic and biotic environmental characteristics. Both lakes types hosted similar fish abundance and biomass, but were substantially different in terms of the fish community structure and species richness. Fish were present in all lakes with at least three species. We discovered a higher α-diversity and a lower β-diversity in managed gravel pit lakes compared to unmanaged lakes. Thus, recreational fisheries management appeared to foster homogenization of fish communities, likely because fisheries managers stock these lakes with desired fish species (e.g., piscivorous fishes and large bodied cyprinids). However, we also detected anthropogenic pathways in the colonization of unmanaged gravel pit lakes, presumably from illegal releases by private people. Importantly, hardly any non-native species were detected in the gravel pits we studied, suggesting that recreational fisheries management not necessarily promotes the spread of exotic species.

**Significance Statement:** Little is known about fish communities in artificially created gravel pit lakes. We compared those managed by recreational fishers with those lacking fisheries management in north-western Germany. We found fishes in all gravel pit lakes and demonstrated a higher α-diversity but more homogenized fish communities in managed gravel pit lakes compared to unmanaged lakes. We did not detect the establishment of relevant abundances of non-natives fishes despite intensive fisheries management.

## 1. Introduction

Freshwater ecosystems have been strongly altered by humans (Dodds *et al.*, 2013). While rivers in the temperate regions have experienced substantial biotic homogenization and habitat loss (Vörösmarty *et al.*, 2010), lakes have mostly suffered from eutrophication, pollution and climate change (Brönmark & Hansson, 2002). Moreover, invasions by non-native species have locally and regionally become an important threat for freshwater ecosystems (Rahel, 2007). Today, freshwater biodiversity is declining at an alarming rate with 37% of Europe’s freshwater fish species categorized as threatened (Freyhof & Brooks, 2011). Habitat loss has been identified as one of the main stressor that impacts biodiversity (Dudgeon *et al.*, 2006; Strayer & Dudgeon, 2010). When properly managed (Lemmens *et al.*, 2013), novel aquatic ecosystems, such as gravel pit lakes (i.e., lentic water bodies created by human use of sand, clay, gravel and other natural resources), reservoirs and ponds, can counteract the freshwater biodiversity crisis by creating secondary habitats for colonization and refuges in the case natural ecosystems deteriorate (Dodson *et al.*, 2000; Santoul *et al.*, 2004, 2009; De Meester *et al.*, 2005; Völkl, 2010; Emmrich *et al.*, 2014; Zhao *et al.*, 2016; Biggs *et al.*, 2017). Gravel pits are often groundwater-fed and not necessarily connected to surrounding river systems (Blanchette & Lund, 2016; Mollema & Antonellini, 2016; Søndergaard *et al.*, 2018); they thus display the interesting biogeographic feature of islands in a landscape (Olden *et al.*, 2010). This characteristic causes a slow colonisation and a potentially low species richness (Magnuson *et al.*, 1998).

In 2014, sand and gravel were extracted from over 28.000 quarries and pits in Europe (UEPG, 2017) resulting in small and isolated gravel pit lakes as common landscape elements in industrialized countries (Blanchette & Lund, 2016; Mollema & Antonellini, 2016; Søndergaard *et al.*, 2018). In Lower Saxony, Germany, more than 37,000 gravel pit lakes smaller than 20 ha exist, and they cover about 70% of all lentic habitats in this region (Manfrin, unpublished data). The fish species richness in natural lakes in northern Germany has been found to be a function of areal size with more species occurring in larger lakes due to higher habitat diversity (Eckmann, 1995). Thus, due to their normally small size below 50 ha, their recent origin after the Second World War within the scope of further industrialization and often isolated location, gravel pit lakes can also be naturally assumed to contain species-poor fish communities, and when newly created may even lack fish populations at all (Scheffer *et al.*, 2006; Søndergaard *et al.*, 2018).

There are several natural pathways for colonization of fishes in gravel pit lakes. When connected with a river fishes can easily colonize these lakes (Molls & Neumann, 1994; Staas & Neumann, 1994; Borcherding *et al.*, 2002). However, the chances of fishes to colonize isolated, recently formed water bodies are low (Scheffer *et al.*, 2006; Strona *et al.*, 2012). Natural colonization is then confined to rare events like massive floods (Pont *et al.*, 1991; Olden *et al.*, 2010) or wind-based dispersal through hurricanes (Bajkov, 1949). Another mean could be passive dispersal of eggs through waterfowl. However, despite frequent claims, this introductory pathways has not been documented with certainty (Hirsch *et al.*, 2018). Thus, it is likely that natural colonization of isolated gravel pit lakes is a very slow process, potentially resulting in species-poor fish communities (i.e., low α-diversity) and high among lake variation in the species pool (i.e., high β-diversity) within a region (Whittaker, 1972; Baselga, 2010).

Human-induced processes like fish introductions, continuous fish stocking or aquarium and bait bucket releases represent an anthropogenic pathways of colonizing human-made freshwater systems (Copp *et al.*, 2010; Gozlan *et al.*, 2010; Olden *et al.*, 2010; Hirsch *et al.*, 2018). In central Europe, by far the majority of gravel pit lakes are managed by recreational anglers organized in clubs and associations (Deadlow *et al.*, 2011). Managers of angling clubs and other fisheries stakeholders regularly engage in fish stocking in lakes and rivers (Cowx, 1994), and this includes gravel pit ecosystems (Arlinghaus, 2006; Arlinghaus *et al.*, 2015; Zhao *et al.*, 2016; Søndergaard *et al.*, 2018). Moreover, illegal releases of garden ponds or aquaria fishes represent the most common pathway of non-native fish dispersal in many areas of the world (Copp *et al.*, 2010; Gozlan *et al.*, 2010; Olden *et al.*, 2010; Patoka *et al.*, 2017) and may thus be widespread in gravel pits as well. Regular stocking may increase α-diversity but reduce β-diversity through the process of biotic homogenization (Radomski & Goeman, 1995; Rahel, 2000, 2002), particularly when fisheries managers stock a common mix of highly desired species (e.g., top predators, Eby *et al.*, 2006). A recent comparison of French gravel pit lakes indeed revealed that the fish community composition was influenced by recreational angling as managed gravel pit lakes hosted more non-native species of high fisheries value, particularly top predators and common carp *Cyprinus carpio* L. compared to unmanaged gravel pit lakes (Zhao *et al.*, 2016).

The objective of the present study was to compare the fish communities between angler-managed and unmanaged gravel pit lakes in north-western Germany. We hypothesized that recreational fisheries management would increase (1) species richness, i.e. α-diversity, (2) the number of piscivorous and other highly desired “game” species and (3) the number of non-native species, such as rainbow trout *Oncorhynchus mykiss* (Walbaum 1792) and topmouth gudgeon *Pseudorasbora parva* (Temminck & Schlegel 1846). We further hypothesized that the lakes managed by anglers host more similar fish communities compared to the unmanged lakes, thereby hypothesizing that (4) recreational fisheries management decreases β-diversity through biotic homogenization.

## 2. Material and Methods

### 2.1 Study lakes and fish sampling

We surveyed the fish communities and a range of limnological lake descriptors in 23 gravel pit lakes located in the lowlands of Lower Saxony, north-western Germany in the Central Plain ecoregion (Fig. 1). For each lake, two ages were determined, the start and the end of gravel mining, as gravel pits start filling up with water and potentially become colonized by fishes already before the end of mining. The depth was measured hydro-acoustically using a Hummingbird Sonar (Type 788ci) in parallel transects spaced about 30 m apart. These data were used to calculate contour maps using ordinary kriging in R (for further details see Supplementary of Monk & Arlinghaus, 2017). The contour maps were used to extract key morphometric variables of the lake (mean depth, maximum depth, shoreline length and area), including estimation of areas covered by different depth strata according to the CEN standard (2015) for the sampling of lake fish communities with multimesh gillnets (0 - 2.9 m, 3 - 5.9 m, 6 – 11.9 m, 12 – 19.9 m and 20 – 34.9 m). These data were also used for the calculation of the shoreline development factor (Osgood, 2005) and the extension of the littoral zone (defined as area between 0 and 2.9 m depth). We mapped macrophytes in summer during full vegetation with a Simrad NSS7 evo2 echosounder with a Lowrance TotalScan Transducer in parallel transects spaced about 30 m apart, similar to the contour maps. Macrophyte coverage and average height were calculated by kriging using a commercial software (Winfield *et al.*, 2015; Valley, 2016; www.gofreemarine.com/biobase/).

**Figure 1:**
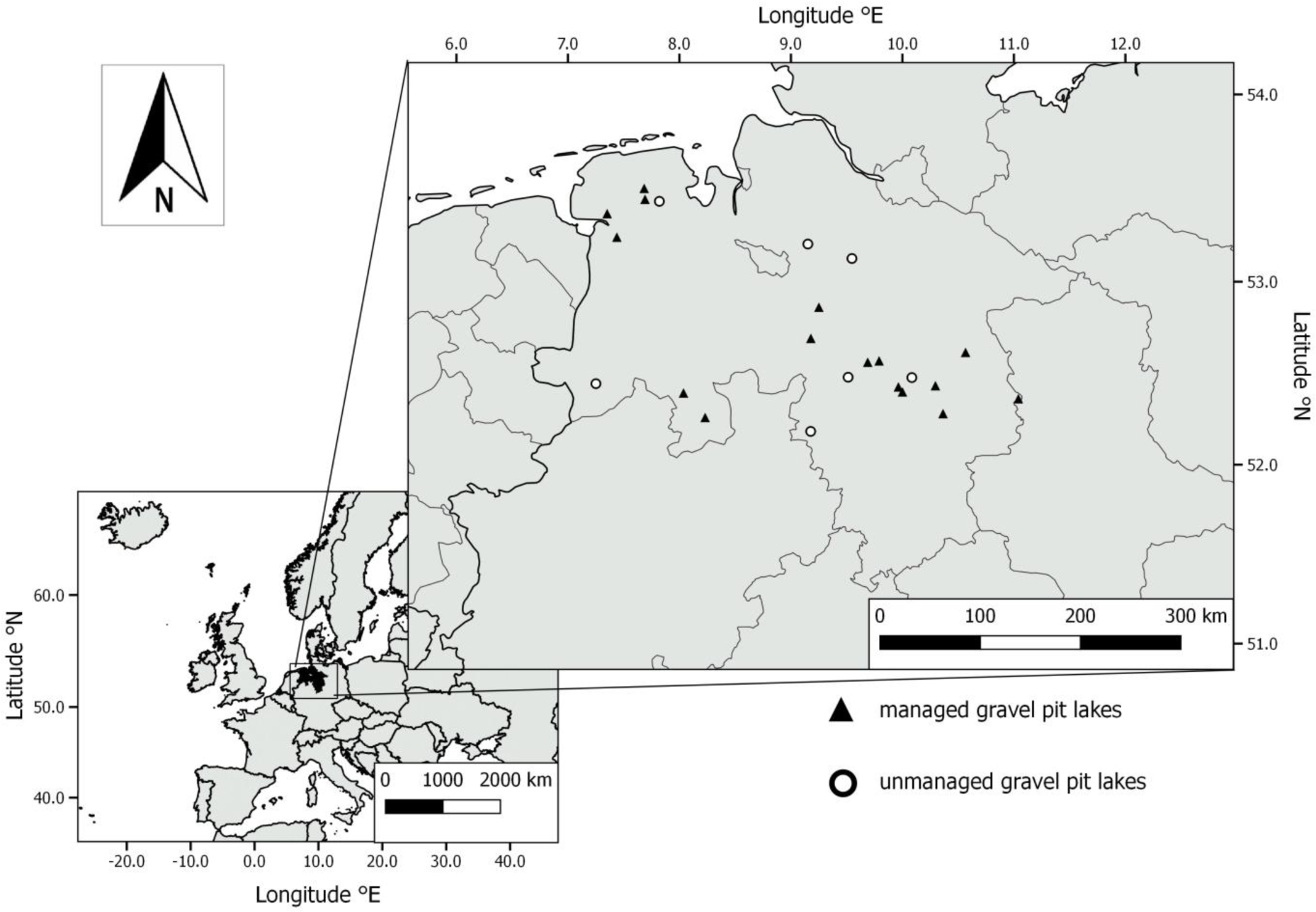
Location of the sampling lakes in Lower Saxony, north-western Germany, Europe

The fish communities were sampled using day-time electrofishing in the littoral and multimesh gillnets in the benthic and profundal zones in autumn 2016 and 2017. During each fish sampling campaign, the lake’s Secchi depth, conductivity and pH-value were measured. In addition, at the deepest point of the lake an oxygen-depth-temperature profile was taken in steps of 50 cm using a WTW Multi 350i sensor, WTW GmbH, Weilheim, Germany, and epilimnic water samples were taking for analyzing total phosphorus concentrations (TP) and chlorophyll a (Chl a). In the laboratory TP was determined using the molybdenum blue method (ISO, 2004; Zwirnmann et al., 1999) and Chl a was determined using high performance liquid chromatograph (Mantoura & Llewellyn, 1983; Wright *et al.*, 1991).

Littoral electrofishing was conducted from a boat by a two person crew using a FEG 8000 electrofishing device (8,0 kW; 150 - 300V / 300 - 600V; EFKO Fischfanggerä te GmbH Leutkirch) with one anodic hand net (40 cm diameter and mesh size 6 mm) and a copper cathode. Prior to sampling the shoreline was divided in transects measuring between 50 and 120 m depending on the available shoreline habitat. Shoreline habitats covered reeds, overhanging trees and branches, submersed and emersed macrophytes, unvegetated littoral zones with no or low terrestrial vegetation (in particular representing angling sites) and mixed habitats that were not dominated by one of these structures. Each transect was sampled separately. The number of transects varied between 4 and 27, depending on the lake size. The length of all transects summed up to the whole lake shore except for the two largest lakes where in total only about two thirds of the shoreline were fished using random selection of transects. Littoral electrofishing was conducted in 16 managed and 4 unmanaged lakes in autumn 2016 (late August to early October when the epilimnion temperature was > 15^°^C) and multimesh gillnets were set overnight for approximately 12 hours following CEN (2015). An additional electrofishing sampling of the entire shoreline was carried out in autumn 2017 (late August to mid-October). Additionally, in autumn 2017 three further unmanaged gravel pit lakes (for a total sample of seven unmanaged lakes) were sampled by littoral electrofishing of the whole shoreline and multimesh gillnets following the same procedure as in 2016. Electrofishing data were standardized by meter shoreline fished for estimation of lake-wide catch per unit effort data as relative abundance index.

The multimesh gillnets differed slightly from the CEN standard (Appleberg, 2000; CEN, 2015) in a way that we used nets with four additional mesh sizes to attempt to also representatively capture large fishes up to 530 mm total length (Šmejkal *et al.*, 2015). The benthic gillnets had a length of 40 m, a height of 1.5 m and were composed of 16 mesh-size panels each being 2.5 m long with mesh sizes of 5, 6.25, 8, 10, 12.5, 15.5, 19.5, 24, 29, 35, 43, 55, 70, 90, 110 and 135 mm. For lakes < 20 ha the European gillnet sampling standard (CEN, 2015) considers a minimum of 8 or 16 gillnets, depending on whether the maximum depth is below or exceeds 12 m, respectively. Using such a fixed minimum number of nets would strongly bias the fishing pressure in the substantially smaller systems that we sampled, i.e., with 8 nets the encounter probabilities of a fish with a net would be much higher in a small system of, say, 1 ha, compared to a larger ecosystem of 20 ha. This would bias the lake-wide CPUE estimates. Therefore, we adapted the number of gillnets set in each lake by applying the minimum number of 16 standard gillnets to the largest lake in our sample (Meitzer See), which is 19.6 ha and has a maximum depth of 23.5 m. These dimensions are close to the smallest lake size of a deep lake in the CEN standard of 20 ha. For this reference lake, we estimated the quotient of area of the 16 gillnets to total lake area as a measure of “gillnet pressure”. Using this target we calculated the appropriate gillnet numbers in smaller lakes to achieve the same “gillnet pressure” in each lake.

The final number of gillnets set in each lake were distributed following a stratified sampling design by depth strata, where number of gillnets by stratum were set in proportion of the share of each depth stratum’s area to total lake surface area (CEN, 2015). Gravel pit lakes with an area larger than 10 ha or a maximum depth of ≥ 10 m were additionally sampled with pelagic multimesh gillnets to record open water species not captured otherwise (CEN, 2015). One pelagic multimesh gillnet was set in each of the following vertical depth strata: 0 - 1.5 m, 3 – 4.5 m, 6 – 7.5 m, 9 - 10.5 m and 12 – 13.5 m, but only if the depth strata contained >1 mg O_2_ L^-1^. Note the pelagic gillnets were only used to complete the species inventory (presence-absence data) as recommended in the CEN standard (CEN, 2015), but not used for the lake wide fish abundance and biomass estimates. Lake-wide biomass and abundance estimates were estimated as stratified means per area and night fished as recommended by CEN (2015).

Total length of the fishes that were captured by electrofishing and gillnetting were measured to the nearest mm and the weight was to the nearest g. In case of higher sample sizes, at least 10 fish per species and 2 cm length class were measured and weighted. Afterwards fishes were only measured and the weight was calculated with length-weight regressions from that specific lake. Only in rare case of catching several hundreds of 0+ fish by electrofishing, a subsample was measured and weighted. Afterwards all the other fish were weighted together and the number and length-frequency distribution of the whole sample was calculated using the length-frequency distribution of the subsample.

### 2.2 Fish community descriptors

For all calculations and analyses, data from 2016 and 2017 were pooled. This results in electrofishing data in 20 lakes from two years and in three lakes from only one year. Furthermore, data from one autumn sampling per lake with multimesh gillnets were analyzed.

Species richness, number of piscivorous species, number of small-bodied non-game fish (after Emmrich *et al.*, 2014), number of threatened species (after the Red List of Lower Saxony (LAVES, 2011), the Red List of Germany (Freyhof, 2009) and the European Red List (Freyhof & Brooks, 2011)) and number of non-native species (after Wiesner *et al.*, 2010 and Wolter & Röhr, 2010) were calculated to describe the lake fish community based on electrofishing (littoral zone) and multimesh gillnet data (benthic and for species richness also the pelagic zone). Perch *Perca fluviatilis* (L.) > 150 mm and eel *Anguilla anguilla* (L.) > 500 mm total length were accounted to the piscivorous fish guild, following Emmrich *et al.* (2014). Cyprinid hybrids were listed as fish caught in the gravel pit lakes (Table 1), but excluded from further analyses of species-specific patterns.

**Table 1:** Common and scientific names, frequency of occurrence and relative frequency in the lake types of the fish species caught in 16 managed and 7 unmanaged gravel pit lakes using littoral electrofishing and benthic and pelagic gillnetting.

| Common name | Scientific name | Frequency of occurrence in managed lakes | Frequency of occurrence in unmanaged lakes | Relative frequency in managed lakes (electrofishing) | Relative frequency in unmanaged lakes (electrofishing) | Relative frequency in managed lakes (gillnetting) | Relative frequency in unmanaged lakes (gillnetting) |
| --- | --- | --- | --- | --- | --- | --- | --- |
| Perch† | *Perca fluviatilis* L. | 100.0 | 28.6 | 42.50 | 19.98 | 62.35 | 27.99 |
| Roach | *Rutilus rutilus* (L.) | 100.0 | 14.3 | 7.05 | 1.19 | 24.31 | 14.11 |
| Tench | *Tinca tinca* (L.) | 93.8 | 28.6 | 3.53 | 3.45 | 0.61 | 0.69 |
| Eel†§ | *Anguilla anguilla* (L.) | 93.8 | 0.0 | 13.24 | 0.00 | 0.00 | 0.00 |
| Pike†§ | *Esox lucius* L. | 87.5 | 14.3 | 3.88 | 5.42 | 0.28 | 0.00 |
| Rudd | *Scardinius erythrophthalmus* (L.) | 68.8 | 42.9 | 14.50 | 16.82 | 1.74 | 16.98 |
| Bream | *Abramis brama* (L.) | 68.8 | 14.3 | 6.77 | 0.00 | 5.21 | 0.17 |
| Carp | *Cyprinus carpio* L. | 56.3 | 42.9 | 0.35 | 0.15 | 0.17 | 1.24 |
| Ruffe‡ | *Gymnocephalus cernua* (L.) | 56.3 | 0.0 | 0.31 | 0.00 | 2.69 | 0.00 |
| Pikeperch† | *Sander lucioperca* (L.) | 50.0 | 0.0 | 0.01 | 0.00 | 0.69 | 0.00 |
| White bream | *Blicca bjoerkna* (L.) | 43.8 | 0.0 | 1.91 | 0.00 | 1.71 | 0.00 |
| Prussian carp | *Carassius gibelio* (Bloch 1782) | 12.5 | 28.6 | 2.67 | 2.83 | 0.23 | 9.14 |
| European catfish†§ | *Silurus glanis* L. | 12.5 | 14.3 | 0.06 | 0.00 | 0.02 | 0.00 |
| Cyprinid hybrid | *Rutilus x Abramis* | 12.5 | 0.0 | 0.00 | 0.00 | 0.00 | 0.00 |
| Topmouth gudgeon‡¶ | *Pseudorasbora parva* (Temminck & Schlegel 1846) | 6.3 | 14.3 | 0.01 | 0.01 | 0.00 | 0.00 |
| Bitterling‡§ | *Rhodeus amarus* (Bloch 1782) | 6.3 | 0.0 | 0.02 | 0.00 | 0.00 | 0.00 |
| European whitefish | *Coregonus lavaretus* (L.) | 6.3 | 0.0 | 0.00 | 0.00 | 0.00 | 0.00 |
| Spined loach‡§ | *Cobitis taenia* L. | 6.3 | 0.0 | 0.39 | 0.00 | 0.00 | 0.00 |
| Bleak‡ | *Alburnus alburnus* (L.) | 6.3 | 0.0 | 2.79 | 0.00 | 0.00 | 0.00 |
| Sunbleak‡ | *Leucaspis delineatus* (Heckel 1843) | 0.0 | 42.9 | 0.00 | 27.51 | 0.00 | 15.18 |
| Nine-spined stickleback‡ | *Pungitius pungitius* (L.) | 0.0 | 42.9 | 0.00 | 21.80 | 0.00 | 6.04 |
| Gudgeon‡ | *Gobio gobio* (L.) | 0.0 | 28.6 | 0.00 | 0.72 | 0.00 | 4.69 |
| Stone loach‡ | *Barbatula barbatula* (L.) | 0.0 | 14.3 | 0.00 | 0.08 | 0.00 | 3.76 |
| Brown bullhead†¶ | *Ameiurus nebulosus* (Lesueur 1819) | 0.0 | 14.3 | 0.00 | 0.05 | 0.00 | 0.00 |

† Piscivorous species (perch > 15 cm total length (TL) and eel > 50 cm TL were classified piscivorous)
‡ Small-bodied non-game fish
§ Threatened species in Lower Saxony
¶ Non-native species

Species richness was used to compare the α-diversity between the lake types. The number of piscivorous species was used as a fish community descriptor as anglers prefer to catch predatory fishes and regularly stock those (Arlinghaus *et al.* 2015). We also assessed the number of small-bodied non-game fish species as many of these species are relevant in a conservation context. Also, many small-bodied species are pioneer colonizer of lakes, e.g. sunbleak *Leucaspius delineates* (Heckel 1843) (Kottelat & Freyhof, 2007). The number of threatened species was contrasted between the two lake types to assess the potential impact of fisheries management on fish conservation objectives. Furthermore, the number of non-native species was compared among lake types, as fish stocking is believed to promote the spread of exotic fishes, particularly in gravel pit lakes (Zhao *et al.*, 2016; Søndergaard *et al.*, 2018).

To compare the relative fish abundance and the abundance-based community descriptors (piscivorous species, small-bodied non-game fish species, threatened species, non-native species) as well as the Shannon diversity index (Shannon, 1948) between the two lake types the mean lake-specific catch per unit effort (CPUE) with individuals per shoreline length (N / 50 m) or gillnet area (N / 100 m^2^) was used (number per unit effort, NPUE). CPUE data regarding the biomass per shoreline length (g / 50 m) or gillnet area (N / 100 m^2^) were also calculated (biomass per unit effort, BPUE). As NPUE data can be affected by the catch of a school of small fishes and BPUE can be affected by the catch of single, large individuals, both calculations were used in the analysis to provide a complete picture. In addition to species numbers, NPUE and BPUE data from electrofishing and gillnetting as well as the Shannon index were used to assess the fish biodiversity and community composition between the two lake types.

### 2.3 Statistical analysis

To test for mean differences between the two lake types regarding the limnological lake characteristics and the biodiversity descriptors of the fish community, a Welch two sample t-test was conducted when raw variables or log_10_-transformed variables were normally distributed and showed homogeneity of variances. In all other cases, a Wilcoxon rank sum test was performed.

Following Anderson *et al.* (2011), the β-diversity of the fish communities in managed and unmanaged gravel pit lakes was visualized by non-metric multidimensional scaling (nMDS; Kruskal, 1964) using Bray-Curtis distances on both species number and abundance data. Afterwards a permutation test for homogeneity of multivariate dispersions (permutations: N=9999) was performed to test for significant differences in the fish community composition. With a similarity percentage analysis (SIMPER; permutations: N=999; Clarke, 1993), the species strongly contributing to the average dissimilarity between the two lake types were identified. All statistical analyses were conducted using R version 3.2.2 (R Core Team, 2016) and the package *vegan* (Oksanen *et al.*, 2018).

## 3. Results

Managed gravel pit lakes varied between 1.0 and 19.6 ha in size with a shoreline length ranging from 415 to 2660 m; unmanaged gravel pit lakes ranged from 2.2 to 11.4 ha in size and varied between 727 and 2060 m in shoreline length. The two lake types did not differ statistically in any morphological variable (Fig. 2): area (Welch two sample t-test, t = 0.728; p = 0.476), shoreline length (Welch two sample t-test, t = 0.706; p = 0.490), shoreline development factor (Wilcoxon rank sum test, W = 53.5; p = 0.867), mean depth (Welch two sample t-test, t = 0.496; p = 0.635), maximum depth (Wilcoxon rank sum test, W = 58; p = 0.922) and share of the littoral (Welch two sample t-test, t = −0.748; p = 0.471). While a difference in lake age in terms of start of mining was detected (managed: 43.4 ± 8.7 a (mean ± SD); unmanaged: 30.4 ± 9.7 a (mean ± SD); Welch two sample t-test, t = 3.03, p = 0.012), no differences were detected for the lake age at the end of mining (managed: mean = 29.4 ± 12.4 a (mean ± SD); unmanaged: mean = 21.6 ± 11.5 a (mean ± SD); Welch two sample t-test, t = 1.475, p = 0.165). Furthermore, no differences among lake types were detected for the variables reflecting lake productivity: total phosphorus (TP; Welch two sample t-test, t = −0.285, p = 0.781), chlorophyll a (Chl a; Welch two sample t-test, t = −1.433, p = 0.181) and Secchi depth (Welch two sample t-test, t = 0.530, p = 0.608). The relatively low mean values of TP and Chl a indicated that the lakes were predominantly mesotrophic. The two lake types also, on average, did not differ in conductivity (Welch two sample t-test, t = 0.903, p = 0.388) and pH-value (Welch two sample t-test, t = −0.920, p = 0.383). Macrophyte data revealed no differences between the lakes types regarding macrophyte coverage (Welch two sample t-test, t = 0.916, p = 0.382), however, the macrophyte hight was larger in managed gravel pit lakes (Welch two sample t-test, t = 2.471, p = 0.036).

**Figure 2:**
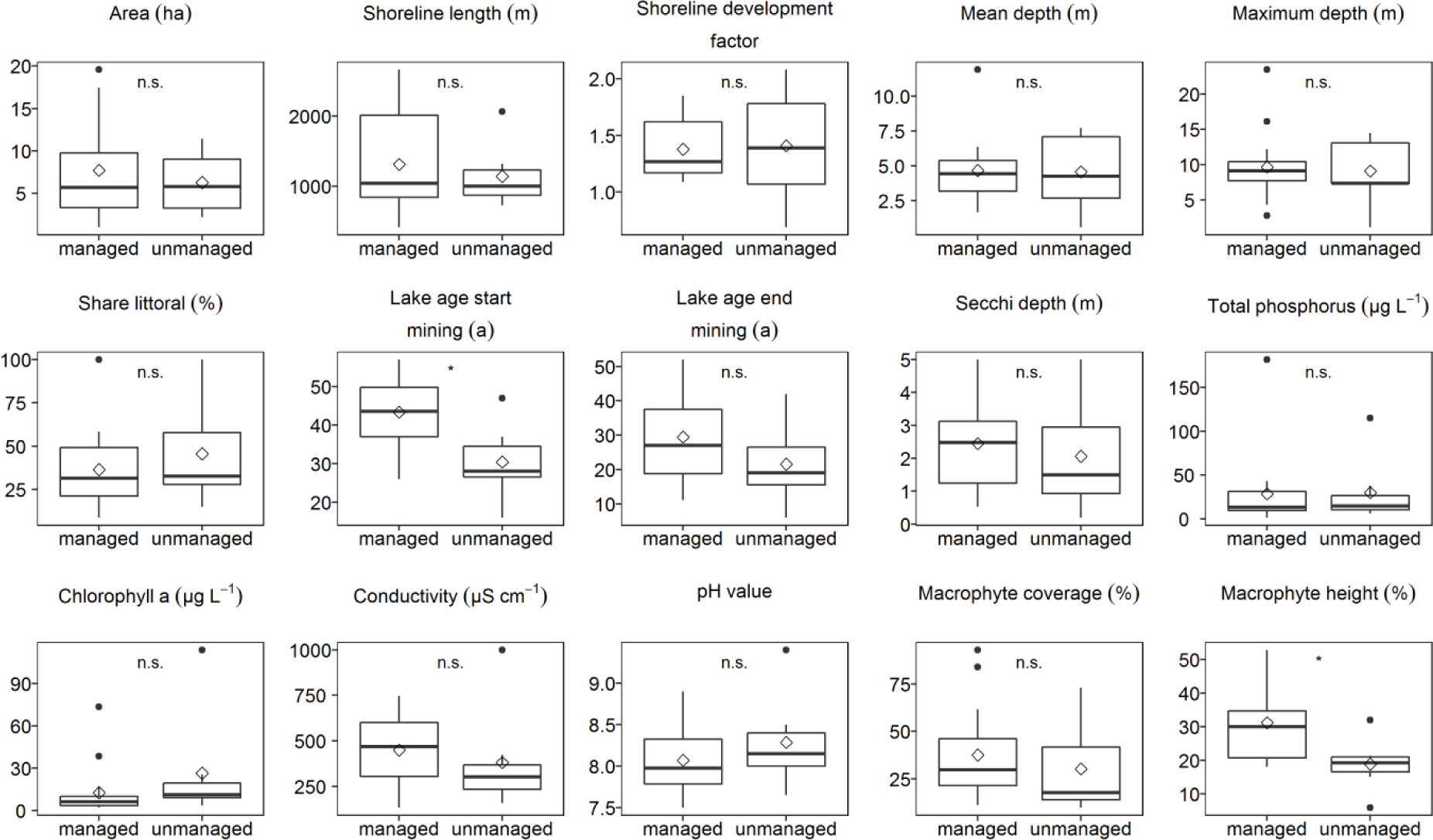
Comparison of the environmental characteristics between managed (N=16) and unmanaged (N=7) gravel pit lakes. The boxes show the 25^th^ to the 75^th^ percentile and the whiskers extent to 1.5 * IQR (inter-quartile range). Median is marked as a solid line, mean as diamond and outliers as circles. Significance levels are * < 0.05; ** < 0.01; *** < 0.001 and n.s. = not significant.

In total 117,214 fishes were sampled, 108,148 individuals by electrofishing and 9,066 by gillnetting. The fish community in the 23 gravel pit lakes consisted of 23 fish species and one hybrid (Table 1). All lakes contained at least three fish species. *Perca fluviatilis* and roach *Rutilus rutilus* (L.) were found in all managed lakes, while they were present in less than a third of the unmanaged lakes. Piscivorous species such as pike *Esox Lucius* L., *Anguilla anguilla* and pikeperch *Sander lucioperca* (L.) were also regularly found in managed, but only occasional or not at all in unmanaged gravel pit lakes (Table 1). Littoral species, such as *Esox lucius*, *Anguilla anguilla* and tench *Tinca tinca* (L.), were mainly or even exclusively caught by electrofishing, while large individuals of less littoral-bound species such as *Perca fluviatilis* and *Rutilus rutilus* as well as *Sander lucioperca* were better detected by gillnetting.

Of the 23 species, *Anguilla anguilla*, *Sander lucioperca*, ruffe *Gynmocephalus cernua* (L.), white bream *Blicca bjoerkna* (L.), bitterling *Rhodeus amarus* (Bloch 1782), European whitefish *Coregonus lavaretus* (L.), spined loach *Cobitis taenia* L. and bleak *Alburnus alburnus* (L.) were only caught in managed gravel pits, while sunbleak *Leucaspinus delineates* (Heckel 1843), nine-spined stickleback *Pungitius pungitius* (L.), gudgeon *Gobio gobio* (L.), stone loach *Barbatula barbatula* (L.) and brown bullhead *Ameiurus nebulosus* (Lesueur 1819) only occurred in unmanaged gravel pits. However, non-native *Ameiurus nebulosus* was only detected as a single individual.

On average, the species richness (Welch two sample t-test, t = 7.61, p < 0.001), number of piscivorous species (Wilcoxon rank sum test, W = 111, p < 0.001) and number of threatened species (Wilcoxon rank sum test, W = 110, p < 0.001) were significantly higher in managed gravel pit lakes compared to unmanaged ones (Fig. 3). No differences between the two lake types were found in the number of small-bodied non-game fish species (Wilcoxon rank sum test, W = 37, p = 0.179) and the number of non-native species (Wilcoxon rank sum test, W = 43.5, p = 0.153). However, in total, only four individual non-native fishes were caught, three specimens of *Pseudorasbora parva* and one specimen of *Ameiurus nebulosus*. The Shannon index revealed an overall greater diversity for the littoral fishes in terms of both abundance and biomass (NPUE and BPUE) and for the whole lake fish community biomass estimate (BPUE) in managed gravel pit lakes compared to unmanaged ones (Table 2).

**Table 2:** Comparison between the two management types for NPUE and BPUE of electrofishing and multimesh gillnet data on the total catch and the catch of the selected fish community descriptors in gravel pit lakes in Germany.

| | | Mean (± S.D.) | | Median (Range) | | p-value |
| --- | --- | --- | --- | --- | --- | --- |
|  |  | Managed lakes | Unmanaged lakes | Managed lakes | Unmanaged lakes |  |
| Littoral estimate<br>(electrofishing) NPUE<br>(N/50m) | Total abundance | 30.4 (±23.7) | 538.4 (±1306.7) | 22.8 (7.3 - 94.7) | 33.6 (3.8 - 3499.6) | 0.368 |
| | Piscivorous fishes | 1.7 (±1.2) | 0.5 (±0.7) | 1.5 (0.2 - 4.2) | 0.0 (0.0 - 1.6) | **0.007** |
| | Small-bodied non-game fishes | 0.4 (±1.6) | 525.4 (±1311.8) | 0.01 (0.0 - 6.5) | 12.7 (0.0 - 3498.5) | **0.031** |
| | Threatened fishes | 2.8 (±1.8) | 0.4 (±0.6) | 2.8 (0.1 - 6.7) | 0.0 (0.0 - 1.4) | **0.001** |
| | Non-native fishes | 0.0 (±0.01) | 0.01 (±0.02) | 0.0 (0.0 - 0.06) | 0.0 (0.0 - 0.1) | 0.189 |
| | Shannon index | 1.1 (±0.4) | 0.5 (±0.3) | 1.1 (0.6 - 1.6) | 0.6 (0.04 - 0.9) | **0.002** |
| Littoral estimate<br>(electrofishing) BPUE<br>(g/50m) | Total abundance | 798.9 (±525.6) | 798.1 (±829.9) | 687.6 (55.9 - 2082.6) | 472.9 (21.5 - 1920.8) | 0.998 |
| | Piscivorous fishes | 451.4 (±406.87) | 94.4 (±152.0) | 318.8 (21.5 - 1556.4) | 0.0 (0.0 - 360.5) | **0.009** |
| | Small-bodied non-game fishes | 0.7 (±1.8) | 216.4 (±523.1) | 0.05 (0.0 - 7.3) | 11.9 (0.0 - 1400.9) | **0.026** |
| | Threatened fishes | 304.3 (±248.8) | 55.9 (±134.8) | 259.9 (25.9 - 973.8) | 0.0 (0.0 - 360.5) | **0.005** |
| | Non-native fishes | 0.01 (±0.03) | 22.3 (±59.1) | 0.0 (0.0 - 0.1) | 0.0 (0.0 - 156.3) | 0.154 |
| | Shannon index | 1.1 (±0.3) | 0.7 (±0.3) | 1.1 (0.4 - 1.6) | 0.7 (0.2 - 1.0) | **0.012** |
| Whole lake estimate<br>(multimesh gillnet)<br>NPUE (N/100m<sup>2</sup>) | Total abundance | 104.5 (±69.6) | 76.9 (±26.0) | 91.8 (23.7 - 235.7) | 77.1 (40.6 - 111.8) | 0.624 |
| | Piscivorous fishes | 6.6 (±4.9) | 1.9 (±3.4) | 6.0 (0.3 - 19.3) | 0.0 (0.0 - 8.2) | **0.009** |
| | Small-bodied non-game fishes | 1.5 (±2.3) | 8.9 (±12.9) | 0.2 (0.0 - 6.6) | 0.0 (0.0 - 33.5) | 0.564 |
| | Threatened fishes | 0.2 (±0.3) | 0.0 (±0.0) | 0.0 (0.0 - 1.2) | 0.0 (0.0 - 0.0) | 0.105 |
| | Non-native fishes | 0.0 (±0.0) | 0.0 (±0.0) | 0.0 (0.0 - 0.0) | 0.0 (0.0 - 0.0) | NA |
| | Shannon index | 0.8 (±0.3) | 0.5 (±0.4) | 0.9 (0.04 - 1.4) | 0.6 (0.07 - 1.0) | 0.098 |
| Whole lake estimate<br>(multimesh gillnet)<br>BPUE (g/100m<sup>2</sup>) | Total abundance | 3,481.1 (±2,016.4) | 3,048.8 (±1,756.0) | 2919.2 (496.2 - 6,999.9) | 3707.7 (98.3 - 4,682.3) | 0.613 |
| | Piscivorous fishes | 948.6 (±771.6) | 327.8 (±694.1) | 700.6 (12.1 - 2,602.3) | 0.0 (0.0 - 1,857.9) | **0.009** |
| | Small-bodied non-game fishes | 12.6 (±22.7) | 29.6 (±39.4) | 1.6 (0.0 - 76.8) | 0.0 (0.0 - 89.8) | 0.773 |
| | Threatened fishes | 53.9 (±142.0) | 0.0 (±0.0) | 142.0 (0.0 - 518.3) | 0.0 (0.0 - 0.0) | 0.105 |
| | Non-native fishes | 0.0 (±0.0) | 0.0 (±0.0) | 0.0 (0.0 - 0.0) | 0.0 (0.0 - 0.0) | NA |
| | Shannon index | 1.1 (±0.3) | 0.5 (±0.4) | 1.2 (0.6 - 1.7) | 0.7 (0.02 - 1.1) | **0.019** |

**Figure 3:**
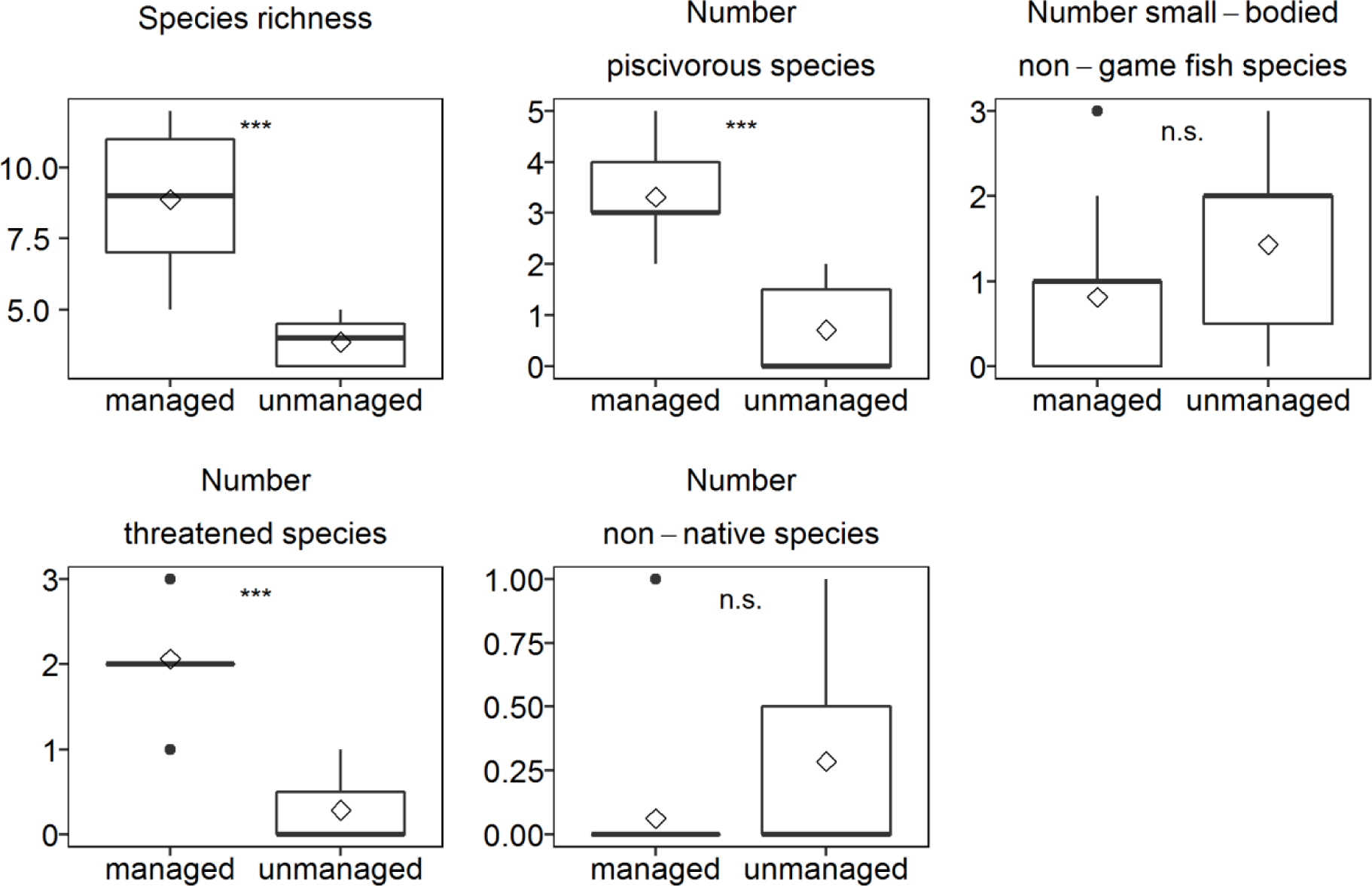
Descriptors of the fish community derived from electrofishing and multimesh gillnetting in managed (N=16) and unmanaged (N=7) gravel pit lakes. The boxplots show the 25^th^ to the 75^th^ percentile (box) and the whiskers extent up to 1.5 * IQR (where IQR is the inter-quartile range). Median is marked as a solid line, mean as diamond and outliers as circles. Significance levels are * < 0.05; ** < 0.01; *** < 0.001 and n.s. = not significant.

Significantly greater abundances (both NPUE and BPUE for both gear types) were found for piscivorous fishes in managed gravel pit lakes compared to unmanaged lakes (Table 2). By contrast, significantly larger abundances and biomasses of small-bodied non-game fishes were detected in unmanaged gravel pit lakes compared to managed ones, but only in the littoral community sampled by electrofishing. For threatened species (*Anguilla anguilla*, *Esox lucius*, European catfish *Silurus glanis* L., *Rhodeus amarus* and *Cobitis taenia*) higher littoral abundances (NPUE and BPUE, respectively) were detected in managed lakes compared to unmanaged lakes. Only four non-native individuals were caught in the littoral by electrofishing and none by multimesh gillnets, meaning that the abundance and biomass of non-natives bordered detectability and accordingly did not differ among lake types.

To investigate differences of the gravel pit fish communities regarding β-diversity, nMDS biplots were constructed by fishing gear using presence-absence data (Appendix) and using abundance and biomass data (NPUE and BPUE; Fig. 4). Permutation tests revealed significantly greater β-diversity for the littoral (NPUE: F = 6.615, p = 0.017; BPUE: F = 11.886, p = 0.002) and benthic fish community (NPUE: F = 13.595, p = 0.001; BPUE: F = 10.106, p = 0.005) in unmanaged gravel pit lakes compared to managed lakes. These differences were revealed by all three means of assessing the fish community (presence-absence, abundance and biomass).

**Fig. 4:**
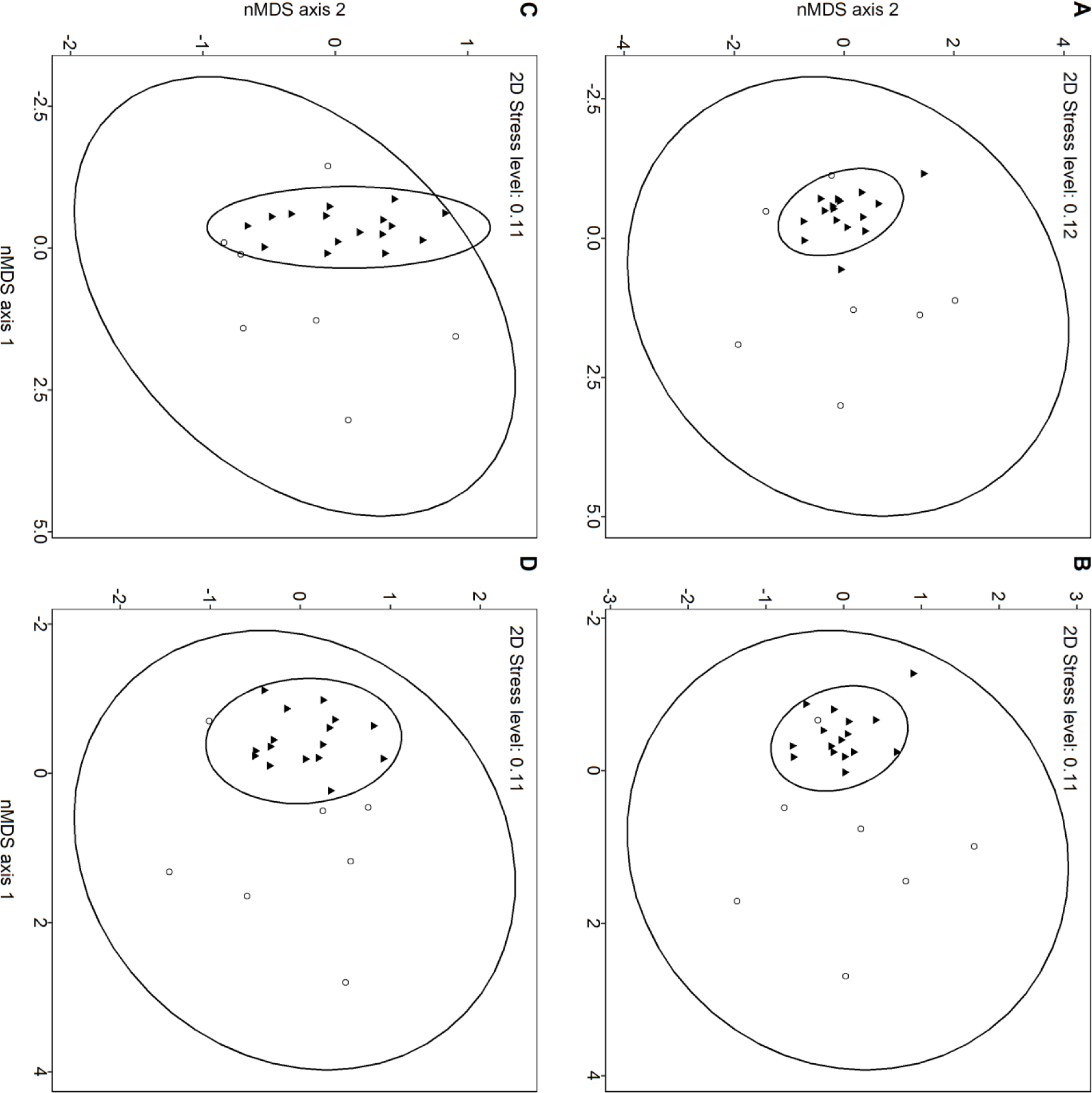
Non-metric multidimensional scaling (nMDS) of the fish community structures with **A** electrofishing NPUE data, **B** electrofishing BPUE data, **C** gillnetting NPUE data and **D** gillnetting BPUE data. Solid triangles represent managed and open circle represent unmanaged gravel pit lakes. The ellipses show the 95% confidence intervals.

*Leucaspinus delineatus*, *Perca fluviatilis*, rudd *Scardinius erythrophthalmus* (L.) and *Pungitius pungitius* contributed 74.6% to the difference between the two lake types in the littoral fish community assessed using electrofishing abundance data (NPUE; Table 3). *Leucaspinus delineatus* and *Pungitius pungitius* were not detected in managed gravel pit lakes, and their contribution to differences in the littoral fish community among lake types was significant (*Leucaspinus delineatus*: p = 0.014, *Pungitius pungitius*: p = 0.013). In terms of littoral fish biomass (electrofishing BPUE data), *Anguilla anguilla*, Prussian carp *Carassius gibelio* (Bloch 1782) and *Esox lucius* contributed most to the difference between the two lake types, but due to high among lake variation in biomass for these species only littoral *Perca fluviatilis* biomass significantly differentiated among managed and unmanaged gravel pit lakes (p = 0.037) revealing significantly greater biomasses in managed lakes.

**Table 3:**
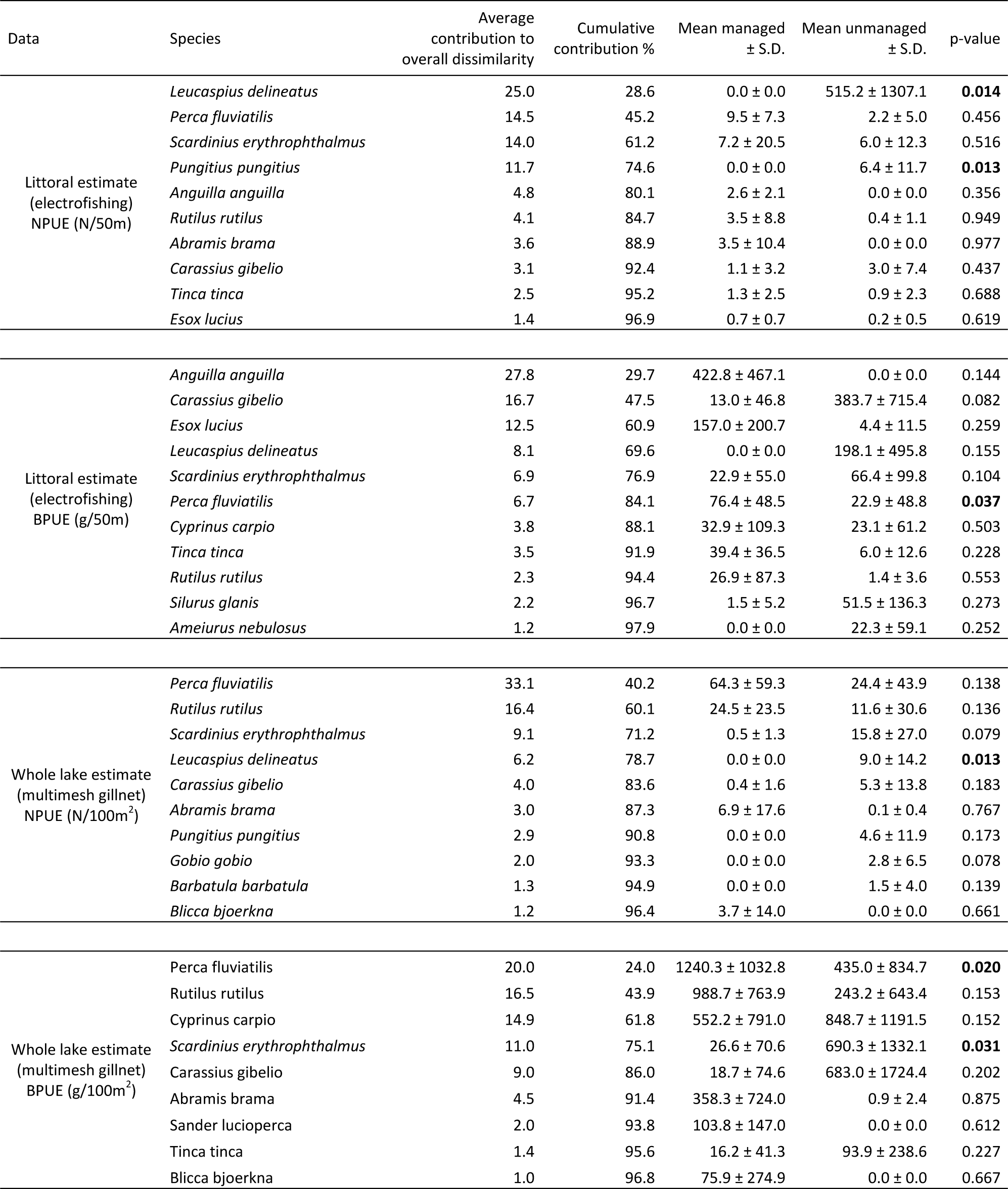
Results of a similarity percentage analysis (SIMPER) for NPUE and BPUE data of electrofishing and multimesh gillnetting, including average dissimilarity, cumulative % contribution to the average dissimilarity, mean and standard deviation for managed and unmanaged gravel pit lakes. Only species explaining more that 1% of the average dissimilarity are presented.

When taking the multimesh gillnet data (NPUE and BPUE) as metrics of whole lake fish community descriptors, *Perca fluviatilis* and *Rutilus rutilus* revealed the highest contribution to the difference in the fish community between the two lake types (significant for *Perca fluviatilis*, p = 0.020 with higher whole-lake biomasses found in managed gravel pit lakes). Furthermore, the whole-lake biomass of *Scardinius erythrophthalmus* differed significantly among lake types, with greater average abundance detected in unmanaged lakes (p = 0.031). In terms of abundance (NPUE), *Leucaspinus delineatus* was a significantly discriminatory species who was only found in multimesh gillnets in unmanaged lakes (p = 0.013).

## 4. Discussion

### 4.1 General findings

We compared the fish communities in angler-managed and unmanaged gravel pit lakes. The results supported three out of four of our initial hypotheses. In particular, species richness (H1) and the number of piscivorous species (H2; e.g., *Esox lucius*, *Sander lucioperca*, *Silurus glanis*, *Perca fluviatilis* and *Anguilla anguilla*) as well as the biomass of piscivorous fishes were significantly higher in managed gravel pit lakes compared to unmanaged lakes. Furthermore, we found a larger number of threatened species and higher littoral abundances and biomasses of threatened fishes in managed gravel pit lakes, while there were no differences in the number of small bodied non-game fish species among lake types. Hence, as hypothesized, managed gravel pit lakes were found to contain a higher α-diversity compared to unmanaged lakes. In contrast to our expectations (H3) the catches of non-native fishes were negligible in both lake types and not significantly greater in managed water bodies as initially assumed. In total four individuals of two species of a total of 117,214 sampled individuals were detected in three different gravel pit lakes. By contrast, the final hypothesis (H4) received substantial support as the species-richer fish communities in managed lakes were more similar to each other than the species-poorer fish communities in unmanaged lakes, suggesting biotic homogenization caused by recreational fisheries management.

### 4.2 Robustness of results to sampling methods

Both groups of gravel pits studied in our work, the ones managed by recreational fishing clubs and the unmanaged lakes, were similar in key environmental characteristics, such as morphology (e.g. lake area) and productivity – factors known in shaping lentic fish communities in the temperate regions (e.g. Persson *et al.*, 1991; Jeppesen *et al.*, 2000; Mehner *et al.*, 2005). This underscores that the fish community differences we report were most likely a result of recreational fisheries management. However, we collected data on the lake age with two different starting points, the start of mining and the end of mining. While the end of mining – a variable used in other studies to determine the gravel pit age (Zhao *et al.*, 2016; Søndergaard *et al.*, 2018) - did not differ between the two lake types, the start of mining differed. Therefore, managed gravel pit lakes had a higher chance to be colonized by chance events due to their older age, which could also have contributed to the larger species richness found in managed compared to unmanaged lakes. The second investigated variable that differed between the lake types was marcophyte hight, however, no differences were detected for macrophyte coverage. As gillnets were set, if possible, in areas without large macrophyte hights and only a low percentage of the electrofished littoral was covered by significant amounts of macrophytes, we assume the influence of the differences in macrophyte hight between the lake types on our data as quite low.

We used electrofishing and multimesh gillnetting to sample the fish community in the gravel pit lakes as adequately as possible as it is known that multiple fishing gears are needed to determine species richness and the habitat-specific abundance in certain habitats of lakes (Barthelmes & Doering, 1996; Diekmann *et al.*, 2005; Jurajda *et al.*, 2009; Achleitner *et al.*, 2012; Menezes *et al.*, 2013; Mueller *et al.*, 2017). Three unmanaged gravel pit lakes were only sampled once in 2017 by electrofishing, while all the other lakes, both managed and unmanaged, were electrofished twice in 2016 and 2017. The lower sampling effort in a subset of the unmanaged lakes likely underestimated the presence of rare species and thus, the average species richness metric in unmanaged lakes might suffer from a negative bias (Lyons, 1992; Angermeier & Smogor, 1995; Paller, 1995). However, as a robustness check, when confining the electrofishing data in all 23 lakes to just one sampling event in one year and comparing the mean species richness of managed and unmanaged lakes, identical results to the ones presented here with our increased sampling effort in 20 lakes were revealed (results not shown). Thus, even if we have underestimated species richness in three of the seven unmanaged lakes that were sampled by electrofishing only once, this bias would not be sufficient to alter our results. The results on the lower species richness in unmanaged lakes in the littoral zone thus appear robust to sampling bias.

Multimesh gillnets were used to sample the fish community of the benthic zone following European standards (CEN, 2015) because the electrofishing is confined to shallow littoral zones. We adapted the gillnet numbers to lake size to equalize fishing pressure across lakes that varied twenty-fold in area. Following Šmejkal *et al.* (2015) we also supplemented the standard mesh sizes in multimesh gillnets by a few larger mesh size panels to sample fish up to 530 mm total length as representatively as possibly, and importantly comparatively across lakes. However, certain species known from previous studies to be present in Lower Saxonian gravel pit lakes (Schälicke *et al.*, 2012) and other angler-managed stagnant water bodies in Germany (Borkmann, 2001), such as the native *Cyprinus carpio* and the non-native Asian carp grass carp *Ctenopharyngodon idella* (Valenciennes 1844), silver carp *Hypophthalmichthys molitrix* (Valenciennes 1844) and bighead carp *Hypophthalmichthys nobilis* (Richardson 1845), have probably been underestimated in their biomass or even completely missed in our design. The reasons are twofold. Some of these large-bodied cyprinids, such as carp, are not very vulnerable to gill-nets and biomasses below 50 kg ha^-1^ are below detectability (Bajer *et al.*, 2016). More importantly, many of these large-bodied species are stocking-reliant and they do not naturally recruit. Hence, these species do not produce individuals vulnerable to the mesh sizes we used, and the fishes over 530 mm are largely invulnerable to the sampling gear we used. It is thus very likely that we missed large-bodied cyprinids and also underestimated the biomass present in large-bodied predators in the managed lakes given the sampling gear we used. The effect on our results is twofold. First, the underestimation of large-bodied cyprinid and predatory species in managed lakes would support our findings as we revealed biotic homogenization through the release of desired fish species and a higher biomass of predators in managed lakes with our design. Second, we might have systematically underestimated the presence and biomass of non-native cyprinids in both lakes types. If in reality large-bodied, non-native cyprinids are only present in managed lakes, our findings on the lack of relevant non-native fishes in managed gravel pit lakes might need to be rethought. Further studies using much longer panels of large mesh sizes are needed to detect large-bodied cyprinids in gravel pit lakes (Schälicke *et al.*, 2012), and we recommend such studies in the future.

### 4.3 Species richness and presence of predators and other “game” species

Species richness and the number of piscivorous species were higher in gravel pit lakes managed for recreational fisheries, supporting our first two hypotheses. In our study, species richness functioned as a surrogate for α-diversity. Supporting our results, a greater α-diversity in lakes managed by and for recreational fisheries has previously been demonstrated for gravel pit lakes in southern France (Zhao *et al.*, 2016) and Minnesota lake fish assemblages (Radomski & Goeman, 1995). Additionally, we also detected a higher Shannon diversity based on the littoral fish abundance and the whole lake biomass estimate in managed gravel pit lakes, further underscoring that managed lakes host larger fish biodiversity than unmanaged gravel pit lakes. Fisheries managers tend to introduce and stock preferentially high trophic level species (Eby *et al.*, 2006; Arlinghaus *et al.*, 2015) and additionally large-bodied cyprinid fishes such as *Cyprinus carpio* and *Tinca tinca* in lakes (Arlinghaus *et al.*, 2015) to meet local angler demands (Arlinghaus & Mehner, 2004; Beardmore *et al.*, 2011; Donaldson *et al.*, 2011; Ensinger *et al.*, 2016). Therefore, as newly created gravel pits are initially fish-free (Schurig, 1972), the documented higher number and higher abundance and biomass of piscivorous species in managed gravel pit lakes is explainable as a result of introductory and maintenance stocking of desired species that eventually establish and self-recruit.

The high-demand species *Anguilla anguilla*, *Esox lucius* and *Perca fluviatilis* were found in all or almost all managed gravel pits, and *Sander lucioperca* in half of the lakes. Although natural colonization of gravel pits by naturally recruiting piscivorous species such as *Esox lucius* and *Perca fluviatilis* is possible, the fact we found *Anguilla anguilla* in almost all managed gravel pit lakes (which all lacked a connection to a river) indicates that stocking must also have played a role. Moreover, it is well known that the angling clubs in the region regularly stock piscivorous fishes such as *Esox lucius* and *Sander lucioperca* (Arlinghaus *et al.*, 2015). These predators were hardly found in unmanaged lakes and similarly no individuals of *Anguilla anguilla* were detected in unmanaged gravel pits at all. The effect of stocking on the presence of species is likely strongest in the early introductory phase when abundant ecological niches are available for colonization. Recent research, however, has shown that once a species is naturally recruiting, stocking with juveniles has no effect on biomass and stock size, e.g., in *Esox lucius* (Johnston *et al.* in press; Hühn *et al.*, 2014). This means that once the initial establishment phase is over, continued angler stocking should particularly affect non-naturally recruiting predatory fishes (Johnston *et al.* in press), in our case *Anguilla anguilla*. Indeed, *Anguilla anguilla* represented one of the major dissimilarities between the two lake types following our SIMPER analyses. A higher relative frequency for *Anguilla anguilla* in gravel pit lakes as result of stocking compared to natural lakes has also been reported previously (Emmrich *et al.*, 2014; Arlinghaus *et al.*, 2016). Given the poor conservation status of eel in nature (e.g. Bark *et al.*, 2007; Dekker, 2016), such stocking events into enclosed water bodies seem questionable.

### 4.4 Small-bodied non-game and threatened species

Small-bodied fishes, such as *Rutilus rutilus*, *Alburnus alburnus* or small *Perca fluviatilis*, are usually less desired by anglers compared to larger-bodied predators (Arlinghaus & Mehner, 2004). However, for all species there are subgroups of anglers that target the species preferentially (Beardmore *et al.*, 2011; Ensinger *et al.*, 2016). Moreover, smaller-bodied cyprinids are considered forage for predators and are therefore also regularly stocked in lentic water bodies in Germany (Arlinghaus *et al.*, 2015). We found some of these species, such as *Rutilus rutilus* and *Perca fluviatilis*, to be present in all managed gravel pits, but they only selectively occurred in a few unmanaged lakes. *Perca fluviatilis* and *Rutilus rutilus* are naturally common in German lakes and have previously been documented to be widespread in lentic water bodies in northern Germany and constitute key element of reference fish communities in lakes (Mehner *et al.*, 2005; Emmrich *et al.*, 2014; Ritterbusch *et al.*, 2014). Although natural colonization is of course possible, it is also likely that widespread small-bodied species were introduced through forage fish stockings or through bait bucket releases in managed water bodies, leading to their common distribution across Lower Saxonian gravel pit lakes in frequencies similar to their distribution in managed natural lakes (Emmrich *et al.*, 2014; Ritterbusch *et al.*, 2014). Therefore, it can be concluded that fisheries management also fosters the establishment and spread of common and naturally widespread percid and cyprinid species.

Small-bodied non-game fishes were also found in both lake types, but the non-game species occurrence strongly differed between managed and unmanaged gravel pit lakes. *Gymnocephalus cernua*, *Rhodeus amarus*, *Cobitis taenia* and *Alburnus alburnus* exclusively occurred in managed lakes, while *Leucaspinus delineates*, *Pungitius pungitius*, *Gobio gobio* and *Barbatula barbatula* were only caught in unmanaged lakes. Furthermore, *Leucaspinus delineates* and *Pungitius pungitius* common to unmanaged lakes strongly contributed to the average dissimilarity between the two lake types. However, at the aggregate level both lakes types hosted the same average number of non-game species. Lake-specific occurrences of specific small-bodied non-game species either represents stochastic effects of natural colonization (e.g., through flooding or influx from nearby creeks and canals) or were additionally caused by stocking efforts of angling clubs. Angling clubs regularly engage in the release of non-game fishes for species conservation purposes, but the volume is small (Arlinghaus *et al.*, 2015) and the activity strongly varies by angling club type (Theis, 2016; Theis *et al.*, 2017). Angling-club specific release of non-game species and stochastic events related to establishment and natural colonization (Copp *et al.*, 2010) can then explain the large variation in species presence of small-bodied non-game species among lakes. The differences in non-game species occurrence in the two lake types also explain the significant differences in the abundances and biomasses of small-bodied non-game fishes among lake types. The difference in abundance and biomass of small-bodied non-game species were only detected in the littoral as multimesh gillnet do not representative catch these small fishes, but also littoral habitats can favour different species (Blaber *et al.*, 1989; Gratwicke & Speight, 2005). Furthermore, fish biomass in lakes is primarily driven by bottom-up effects (e.g. Hanson & Leggett, 1982; Lemmens *et al.*, 2018; Matsuzaki *et al.*, 2018) – a finding also revealed in our study at the aggregate biomass level, which did not differ among lake types despite radically different fish community composition. As species richness was substantially lower in unmanaged lakes, the lake type-specific small-bodied non-game species that colonized unmanaged lakes can reach higher biomasses and abundances in these lakes types compared to managed lakes. It may also be possible that the small-bodied non-game species in managed lakes may suffer from competitive bottlenecks caused by competition for zooplankton with small-bodied “game” cyprinids and be affected by predation through greater biomasses of piscivorous fish in these lakes, thereby reducing their biomass in managed compared to unmanaged lakes. However, it is similarly plausible that specific non-game species detected in unmanaged lakes, but not occurring in managed lakes, may never have colonized these lakes due to chance events.

The studied lakes hosted a total number of five regionally threatened species, indicating their potential as biodiversity reservoir (Emmrich *et al.*, 2014). *Anguilla anguilla*, *Rhodeus amarus* and *Cobitis taenia* occurred exclusively in managed lakes, while *Esox lucius* and *Silurus glanis* were caught in both lake types. Note that none of these regionally threated species are listed in the German Red List of freshwater fishes, yet the *Anguilla anguilla* is today globally threatened according to IUCN criteria (Freyhof & Brooks, 2011). The number of regionally threatened species (as judged by their presence on the regional Red List of Lower Saxony) and the littoral abundance and biomasses were significantly higher in managed lakes compared to unmanaged lakes, thereby suggesting managed lakes can function as secondary habitat for hosting threatened species. In particular *Esox lucius* is worth highlighting, as it was caught in 87.5% of the managed, but only in 14.3% of the unmanaged gravel pit lakes. Fisheries management thus can foster the fish conservation value of managed gravel pit lakes as *Esox lucius* typically establishes self-reproducing population in gravel pit lakes after first introduction (Schälicke *et al.*, 2012). With regards to eel, as mentioned above, the continued stocking in enclosed water bodies seems problematic from a conservation standpoint as they do not enlarge the spawning stock biomass.

### 4.5 Presence of non-native fishes

The hypothesized support of non-native species introductions and accumulation of exotics by recreational fisheries management as revealed in a French gravel pit study by Zhao *et al.* (2016) was not confirmed for artificial lakes in north-western Germany. An examination of the French study revealed that the non-native species listed there encompassed many species being native in Germany, but not in France, such as *Cyprinus carpio*, *Sander lucioperca* and *Silurus glanis*. In our study, only two individuals of non-native *Pseudorasbora parva* were found in one of 16 managed lakes, which was most likely unwillingly introduced through poorly sorted stocking of pond-reared *Cyprinus carpio* or poorly sorted wild stocking of cyprinids (e.g. Copp *et al.*, 2005b; Wiesner *et al.*, 2010). In comparison, in two out of seven unmanaged lakes, one individual of each of two non-native species, namely *Pseudorasbora parva* and *Ameiurus nebulosus*, were detected, showing that also unmanaged lakes receive propagule pressure by non-natives. Moreover, in a further unmanaged gravel pit lake many individuals of a golden variant of *Scardinius erythrophthalmus*, common as ornamental fish, were found. Illegal stocking (e.g. release of fish by owners of garden ponds or local anglers interested in establishing desired species in a region) has been shown to contribute as vector for fish dispersal around the globe (Johnson *et al.*, 2009; Hirsch *et al.*, 2018), e.g. illegal goldfish *Carassius auratus* (L.) stocking in Great Britain (Copp *et al.*, 2005a). This is further evidence that the unmanaged lakes in our study were affected by illegal release of fishes by private people. Illegal release, rather than organized fisheries management by fishing clubs, is today believed to constitute the most important pathway for the transfer of non-natives fishes across the world (Copp *et al.*, 2010). One reason for this is that release of non-native fishes is banned in Germany based on Nature Conservation Law and most fisheries are private property and run by trained fisheries managers, many of which are alert of the need to constrain establishment of non-native fishes and rarely engage in stocking non-natives (Arlinghaus *et al.*, 2015; Riepe *et al.*, 2017). While such conditions still allow dispersal of fishes by private anglers illegally (Johnson *et al.*, 2009), even if such activities occurred in our lakes, they have left a limited legacy in the study region. This finding agrees with Wolter & Röhr (2010) who reported that non-native fishes rarely have become invasive in Germany, with river populations of gobies being an exception. We conclude that proper recreational fisheries management is not per se a vector for non-native species establishment and that not managing lakes is not a guarantee for the lack of establishment either.

### 4.6 Biotic homogenization caused by fisheries management

We found in agreement with our expectation that recreational fisheries management contributed to the homogenization of fish faunas, reducing the β-diversity in fish communities compared to unmanaged lakes. Homogenization of fish communities as a result of a range anthropogenic influences (e.g., stocking, urbanization, habitat simplification) has been repeatedly found across the world (Radomski & Goeman, 1995; Rahel, 2000; Villéger *et al.*, 2011). Our work shows that gravel pit lakes in north-western Germany are no exception. But in contrast to other studies, we can exclude non-fishing related impacts to have exerted substantial impacts on our results as the environment of the lakes we studied was rather similar and only the presence or absence of fisheries management discriminated among the lakes. Biotic homogenization not only entails that fish communities become increasingly similar across ecoregions and among ecosystems but that many ecosystems increasingly host environmentally tolerant or species of high fisheries value, thereby increasing the α-diversity locally, but decreasing β-diversity regionally. This result reflects that management based on introductory stocking or maintenance stocking during the management operations of a lake leads to the accumulation of a certain set of desired species in gravel pit lakes that then establish and become self-reproducing (Emmrich *et al.*, 2014) and collectively contribute to a homogenized community that reduces β-diversity.

Homogenization in lentic fish communities over longer periods of time (e.g., 10.000 years after the last glaciation) may also be a natural phenomenon as evidenced by fish community assessments of natural lakes across Europe and in northern Germany who have shown that only a few key environmental gradients (e.g., lake depth and productivity) discriminate among different lentic fish communities (Diekmann *et al.*, 2005; Mehner *et al.*, 2005; Brucet *et al.*, 2013; Ritterbusch *et al.*, 2014). Put differently, lakes having similar limnological characteristics over time also host very similar (i.e., homogenous) fish communities, supporting the results of our managed lakes. One limitation to this statement is that also most of the natural lakes assessed by Diekmann *et al.* (2005), Mehner *et al.* (2005) and Emmrich *et al.* (2014) and used by Ritterbusch *et al.* (2014) to derive reference fish communities were managed for fisheries presently or in the past. In that sense the newly created, yet unmanaged gravel pit lakes with their lake-specific species poor communities do not find any natural parallel in the existing studies. Therefore, we conclude that high β-diversity maybe the natural conditions at least in the initial phases of the succession in lakes. Similar to islands, the chances of fishes to naturally colonize isolated freshwater habitats are low (Scheffer *et al.*, 2006; Strona *et al.*, 2012) and often limited to rare events like massive floods (Pont *et al.*, 1991; Olden *et al.*, 2010), fish rain (Bajkov, 1949) and less likely bird-based dispersal (Riehl, 1991; Hirsch *et al.*, 2018; Martin & Turner, 2018). A further colonization mechanism identified in our work encompasses uncontrolled anthropogenic events, such aquarium, garden pond and bait bucket releases (Padilla & Williams, 2004; Copp *et al.*, 2005a; Hirsch *et al.*, 2018). Overall, also in light to the existing literature it seems that the high β-diversity we found is special to young lake ecosystems in a pioneer status. Importantly, however, as mentioned before, none of the unmanaged lakes were fish-free and they contained at least three fish species, while often lacking piscivorous fish species. Thus, one key message of our work is that also lakes not managed by and for anglers will be colonized by fishes, through both natural and human-assisted means. Our study differs from a recent Danish gravel pit study where the authors reported fish-free systems, but the gravel pit lakes were on average much younger than the ones in our study (Søndergaard *et al.*, 2018).

### 4.7 Conclusions and implications

Our findings offer two conclusions that vary depending on which perspective is taken. First, ongoing homogenization of fish faunas globally (Rahel, 2002; Villéger *et al.*, 2011) can contribute to the freshwater biodiversity crisis (Dudgeon *et al.*, 2006) by reducing the variation in lake fish communities among ecosystems. If such perspective is taken, recreational fisheries management is one contributing factor to homogenization, which maybe judged as undesirable by some. Alternatively and secondly, one has to realize that completely unmanaged lakes and angler-managed lakes will be affected by similar drivers (e.g., introductory or illegal stocking). In a highly urbanized environment it is an illusion to assume natural development of newly created aquatic ecosystems without human impact is possible. If one then acknowledges, based on our work, that proper management of recreational fisheries does not necessarily lead to the development of artificial fish communities with many non-native fishes, using newly created lakes for fisheries can be considered of conservation value despite homogenization of fish communities being largely inevitable. Importantly, recreational fisheries management promotes the rapid establishment of fish communities in new ecosystems that largely resemble similarly structured, managed natural lake ecosystems (Emmrich *et al.*, 2014; Ritterbusch *et al.*, 2014). If new aquatic ecosystems would not be managed, the process of establishment of fishes would likely take substantially longer and would strongly be influenced by stochastic events.

Importantly, most standing waters in Germany are managed under fishing rights laws. Under such conditions, recreational fisheries management contributes to fulfilling state-specific fisheries objectives as specified in the laws, which demand fisheries stakeholders to help establish and maintain near-natural fish communities that match those to be expected under prevailing environmental conditions in natural ecosystems (Arlinghaus, 2017). In all German states except Schleswig-Holstein, gravel pit lakes are legally under the same demand as natural lakes as regards to the goals of fisheries management. However, for artificial waters like gravel pit lakes no reference fish community yet exists, and it is likely that this reference community is species poorer than those documented in natural, managed lakes in Germany (compare Emmrich *et al.* (2014) and Ritterbusch *et al.* (2014)). In the absence of this information, fisheries operators could use fish communities present in environmentally similar (in terms of depth, nutrient content, visibility, habitat structure) natural ecosystems as benchmarks for the species pool expected to be developed in in gravel pit lakes (Arlinghaus *et al.*, 2016). An alternative view maybe that the natural condition of a gravel pit lake is initially fish-free, with slow accumulation of naturally colonized species. While this perspective is valid, our work clearly showed that fish-free lakes are not necessarily to be expected and that anthropogenic introductions also happen in lakes that are not managed by recreational fisheries. Importantly, the number of threatened species in managed and unmanaged lakes did not differ and non-native fishes were rare, implicating that recreational fisheries management is not necessarily a vector for the establishment of artificial fish communities that lack species of conservation value.

## Acknowledgements

We would like to thank all the people participating in the fieldwork, namely Alexander Türck, Leander Höhne, Jara Niebuhr, Philipp Czapla, Andreas Maday, Adrian Schörghöfer, Jasper Münnich, Baiba Prūse and Laura Mehner. Moreover, we thank Angelsportverein Leer u. Umgebung e.V., Bezirksfischereiverband für Ostfriesland e.V., Angler-Verein Nienburg e.V., ASV Neustadt am Rübenberge e.V., Fischereiverein Hannover e.V., Niedersächsisch-Westfälische Anglervereinigung e.V., Stiftung Naturschutz im Landkreis Rotenburg (Wümme), Henning Scherfeld, FV Peine-Ilsede u. Umgebung e.V., SFV Helmstedt u. Umgebung e.V., Verein der Sportfischer Verden (Aller) e.V., Verein für Fischerei und Gewässerschutz Schönewörde u. Umgebung e.V., Steffen Göckemeyer, Thomas Reimer, Michael Wintering and the Angling Association of Lower Saxony for participating in this study. Big thanks go to Robert Morgenstern and Christopher Monk for their help with the contour maps and Miquel Palmer for help with the statistical analysis. Additionally, we are thankful to Thomas Mehner and all the participants of the seminar “Scientific writing” for helpful discussions on an early draft of the manuscript. The study was financed by the German Federal Ministry of Education and Research (BMBF) and the Federal Agency for Nature Conservation (BfN) within the BAGGERSEE-Projekt (grant number: 01LC1320A; www.baggersee-forschung.de).

## Contributions

SM: ideas, data generation, data analysis, manuscript preparation

ME: ideas, data generation, data analysis, manuscript editing

TK: data generation, manuscript editing, funding

CW: manuscript editing, funding

NW: ideas, data generation, data analysis

RA: ideas, data generation, manuscript editing, funding

## References

Achleitner, D., Gassner, H. & Luger, M. (2012) Comparison of three standardised fish sampling methods in 14 alpine lakes in Austria. Fisheries Management and Ecology 19, 352–361 doi.org/10.1111/j.1365-2400.2012.00851.x

Anderson, M. J., Crist, T. O., Chase, J. M., Vellend, M., Inouye, B. D., Freestone, A. L., Sanders, N. J., Cornell, H. V., Comita, L. S., Davies, K. F., et al. (2011) Navigating the multiple meanings of β diversity: A roadmap for the practicing ecologist. Ecology Letters 14, 19–28 doi.org/10.1111/j.1461-0248.2010.01552.x

Angermeier, P. L. & Smogor, R. A. (1995) Estimating number of species and relative abundances in stream-fish communities: effects of sampling effort and discontinuous spatial distributions. Canadian Journal of Fisheries and Aquatic Sciences 52, 936–949 doi.org/10.1139/f95-093

Appleberg, M. (2000) Swedish standard methods for sampling freshwater fish with multi-meshed gillnets. Fiskeriverket Information. 2000, 3–32

Arlinghaus, R. (2006) Overcoming human obstacles to conservation of recreational fishery resources, with emphasis on central Europe. Environmental Conservation 33, 46–59 doi.org/10.1017/S0376892906002700

Arlinghaus, R. (2017) Nachhaltiges Management von Angelgewässern?: Ein Praxisleitfaden. Berichte des IGB, Band 30, 231 S.

Arlinghaus, R. & Mehner, T. (2004) A management-orientated comparative analysis of urban and rural anglers living in a metropolis (Berlin, Germany). Environmental Management 33, 331–344 doi.org/10.1007/s00267-004-0025-x

Arlinghaus, R., Cyrus, E. M., Eschbach, E., Fujitani, M., Hühn, D., Johnston, F., Pagel, T. & Riepe, C. (2015) Hand in Hand für eine nachhaltige Angelfischerei. Ergebnisse und Empfehlungen aus fünf Jahren praxisorientierter Forschung zu Fischbesatz und seinen Alternativen. Berlin: Leibniz-Institut für Gewässerökologie und Binnenfischerei (IGB).

Arlinghaus, R., Emmrich, M., Hühn, D., Schälike, S., Lewin, W.-C., Pagel, T., Klefoth, T. & Rapp, T. (2016) Ufergebundene Fischartenvielfalt fischereilich gehegter Baggerseen im Vergleich zu eiszeitlich entstandenen Naturseen in Norddeutschland. Fischer & Teichwirt 68, 288–291

Bajer, P. G., Beck, M. W., Cross, T. K., Koch, J. D., Bartodziej, W. M. & Sorensen, P. W. (2016) Biological invasion by a benthivorous fish reduced the cover and species richness of aquatic plants in most lakes of a large North American ecoregion. Global Change Biology 22, 3937–3947 doi.org/10.1111/gcb.13377

Bajkov, A. D. (1949) Do fish fall from the Sky? Science 109, 402 doi.org/10.1126/science.109.2834.402

Bark, A., Williams, B. & Knights, B. (2007) Current status and temporal trends in stocks of European eel in England and Wales. ICES Journal of Marine Science 64, 1368–1378 doi.org/10.1093/icesjms/fsm117

Barthelmes, D. & Doering, P. (1996) Sampling Efficiency of Different Fishing Gear used for Fish Faunistic Surveys in stagnant Water Bodies. Limnologica 26, 191–198

Baselga, A. (2010) Partitioning the turnover and nestedness components of beta diversity. Global Ecology and Biogeography 19, 134–143 doi.org/10.1111/j.1466-8238.2009.00490.x

Beardmore, B., Haider, W., Hunt, L. M. & Arlinghaus, R. (2011) The importance of trip context for determining primary angler motivations: Are more specialized anglers more catch-oriented than previously believed? North American Journal of Fisheries Management 31, 861–879 doi.org/10.1080/02755947.2011.629855

Biggs, J., Fumetti, S. & Kelly-Quinn, M. (2017) The importance of small water bodies for biodiversity and ecosystem services: implications for policy makers. Hydrobiologia 793 doi.org/10.1007/s10750-016-3007-0

Blaber, S. J. M., Brewer, D. T. & Salini, J. P. (1989) Species composition and biomasses of fishes in different habitats of a tropical Northern Australian estuary: Their occurrence in the adjoining sea and estuarine dependence. Estuarine, Coastal and Shelf Science 29, 509–531 doi.org/10.1016/0272-7714(89)90008-5

Blanchette, M. L. & Lund, M. A. (2016) Pit lakes are a global legacy of mining: an integrated approach to achieving sustainable ecosystems and value for communities. Current Opinion in Environmental Sustainability 23, 28–34 doi.org/10.1016/j.cosust.2016.11.012

Borcherding, J., Bauerfeld, M., Hintzen, D. & Neumann, D. (2002) Lateral migrations of fishes between floodplain lakes and their drainage channels at the Lower Rhine: Diel and seasonal aspects. Journal of Fish Biology 61, 1154–1170 doi.org/10.1111/j.1095-8649.2002.tb02462.x

Borkmann, I. (2001) Fischereiliche Bonitierung von Still-und Fließgewässern des DAV und des VDSF in Sachsen-Anhalt. In: Weniger, U. (Ed.), Tagungsband zum 4. Landesfischereitag des Landesfischereiverbandes Sachsen-Anhalt e.V. am 17. März 2001 in Blankenburg/Harz. Landesfischereiverband Sachsen-Anhalt e.V., Halle, pp. 17–32.

Brönmark, C. & Hansson, L.-A. (2002) Environmental issues in lakes and ponds: current state and perspectives. Environmental Conservation 29, 290–306 doi.org/10.1017/S0376892902000218

Brucet, S., Pédron, S., Mehner, T., Lauridsen, T. L., Argillier, C., Winfield, I. J., Volta, P., Emmrich, M., Hesthagen, T., Holmgren, K., et al. (2013) Fish diversity in European lakes: Geographical factors dominate over anthropogenic pressures. Freshwater Biology 58, 1779–1793 doi.org/10.1111/fwb.12167

CEN (2015) Water quality—Sampling of fish with multi-mesh gillnets. European Committee for Standardization, EN 14757:2015

Clarke, K. R. (1993) Non-parametric multivariate analyses of changes in community structure. Australian Journal of Ecology 18, 117–143 doi.org/10.1111/j.1442-9993.1993.tb00438.x

Copp, G. H., Wesley, K. J. & Vilizzi, L. (2005a) Pathways of ornamental and aquarium fish introductions into urban ponds of Epping Forest (London, England): The human vector. Journal of Applied Ichthyology 21, 263–274 doi.org/10.1111/j.1439-0426.2005.00673.x

Copp, G. H., Bianco, P. G., Bogutskaya, N. G., Erös, T., Falka, I., Ferreira, M. T., Fox, M. G., Freyhof, J., Gozlan, R. E., Grabowska, J., et al. (2005b) To be, or not to be, a non-native freshwater fish? Journal of Applied Ichthyology 21, 242–262

Copp, G. H., Vilizzi, L. & Gozlan, R. E. (2010) The demography of introduction pathways, propagule pressure and occurrences of non-native freshwater fish in England. Aquatic Conservation: Marine and Freshwater Ecosystems 20, 595–601 doi.org/10.1002/aqc.1129

Cowx, I. G. (1994) Stocking strategies. Fisheries Management and Ecology 1, 15–30 doi.org/10.1111/j.1365-2400.1970.tb00003.x

Deadlow, K., Beard, T. D. & Arlinghaus, R. (2011) A Property Rights-Based View on Management of Inland Recreational Fisheries: Contrasting Common and Public Fishing Rights Regimes in Germany and the United States. American Fisheries Society Symposium 75, 13–38

Dekker, W. (2016) Management of the eel is slipping through our hands Distribute control and orchestrate national protection. ICES Journal of Marine Science: Journal du Conseil 73, 2442–2452 doi.org/10.1093/icesjms/fsw094

Diekmann, M., Brämick, U., Lemcke, R. & Mehner, T. (2005) Habitat-specific fishing revealed distinct indicator species in German lowland lake fish communities. Journal of Applied Ecology 42, 901–909 doi.org/10.1111/j.1365-2664.2005.01068.x

Dodds, W. K., Perkin, J. S. & Gerken, J. E. (2013) Human impact on freshwater ecosystem services: A global perspective. Environmental Science and Technology 47, 9061–9068 doi.org/10.1021/es4021052

Dodson, S. I., Arnott, S. E. & Cottingham, K. L. (2000) The Relationship in Lake Communities Between Primary Productivity and Species Richness. Ecology 81, 2662–2679 doi.org/10.1890/0012-9658(2000)081[2662:TRILCB]2.0.CO;2

Donaldson, M. R., O’Connor, C. M., Thompson, L. A., Gingerich, A. J., Danylchuk, S. E., Duplain, R. R. & Cooke, S. J. (2011) Contrasting Global Game Fish and Non-Game Fish Species. Fisheries 36, 385–397 doi.org/10.1080/03632415.2011.597672

Dudgeon, D., Arthington, A. H., Gessner, M. O., Kawabata, Z.-I., Knowler, D. J., Lévêque, C., Naiman, R. J., Prieur-Richard, A.-H., Soto, D., Stiassny, M. L. J., et al. (2006) Freshwater biodiversity: importance, threats, status and conservation challenges. Biological Reviews of the Cambridge Philosophical Society 81, 163–182 doi.org/10.1017/S1464793105006950

Eby, L. A., Roach, W. J., Crowder, L. B. & Stanford, J. A. (2006) Effects of stocking-up freshwater food webs. Trends in Ecology and Evolution 21, 576–584 doi.org/10.1016/j.tree.2006.06.016

Eckmann, R. (1995) Fish species richness in lakes of the northeastern lowlands in Germany. Ecology of Freshwater Fish 4, 62–69 doi.org/10.1111/j.1600-0633.1995.tb00118.x

Emmrich, M., Schälicke, S., Hühn, D., Lewin, C. & Arlinghaus, R. (2014) No differences between littoral fish community structure of small natural and gravel pit lakes in the northern German lowlands. Limnologica 46, 84–93 doi.org/10.1016/j.limno.2013.12.005

Ensinger, J., Brämick, U., Fladung, E., Dorow, M. & Arlinghaus, R. (2016) Schriften des Instituts für Binnenfischerei e. V. Potsdam-Sacrow. Vol. 44

Freyhof, J. (2009) Rote Liste der im Süßwasser reproduzierenden Neunaugen und Fische (Cyclostomata & Pisces). Naturschutz und Biologische Vielfalt 70, 291–316

Freyhof, J. & Brooks, E. (2011) European Red List of Freshwater Fishes. Luxembourg: Publications Office of the European Union.

Gozlan, R. E., Britton, J. R., Cowx, I. & Copp, G. H. (2010) Current knowledge on non-native freshwater fish introductions. Journal of Fish Biology 76, 751–786 doi.org/10.1111/j.1095-8649.2010.02566.x

Gratwicke, B. & Speight, M. R. (2005) The relationship between fish species richness, abundance and habitat complexity in a range of shallow tropical marine habitats. Journal of Fish Biology 66, 650–667 doi.org/10.1111/j.1095-8649.2005.00629.x

Hanson, J. M. & Leggett, W. C. (1982) Empirical Prediction of Fish Biomass and Yield. Canadian Journal of Fisheries and Aquatic Sciences 39, 257–263 doi.org/10.1139/f82-036

Hirsch, P. E., N’Guyen, A., Muller, R., Adrian-Kalchhauser, I. & Burkhardt-Holm, P. (2018) Colonizing Islands of water on dry land-on the passive dispersal of fish eggs by birds. Fish and Fisheries 1–9 doi.org/10.1111/faf.12270

Hühn, D., Lübke, K., Skov, C., Arlinghaus, R. & Taylor, E. (2014) Natural recruitment, density-dependent juvenile survival, and the potential for additive effects of stock enhancement: an experimental evaluation of stocking northern pike (*Esox lucius*) fry. Canadian Journal of Fisheries and Aquatic Sciences 71, 1508–1519 doi.org/10.1139/cjfas-2013-0636

ISO (2004) Water quality - Determination of phosphorus - Ammonium molybdat spectrometric method. 6878:2004

Jeppesen, E., Jensen, J. P., Søndergaard, M., Lauridsen, T. & Landkildehus, F. (2000) Trophic structure, species richness and diversity in Danish lakes: changes along a phosphorus gradient. Freshwater Biology 45, 201–218 doi.org/10.1046/j.1365-2427.2000.00675.x

Johnson, B. M., Arlinghaus, R. & Martinez, P. J. (2009) Are We Doing All We Can to Stem the Tide of Illegal Fish Stocking? Fisheries 34, 389–394 doi.org/10.1577/1548-8446-34.8.389

Johnston, F., Allen, M., Beardmore, B., Riepe, C., Pagel, T., Hühn, D. & Arlinghaus, R. (2017) How ecological processes shape the outcomes of stock enhancement and harvest regulations in recreational fisheries. Ecological Applications in press doi.org/10.1002/eap.1793

Jurajda, P., Janáč, M., White, S. M. & Ondračková, M. (2009) Small - but not easy: Evaluation of sampling methods in floodplain lakes including whole-lake sampling. Fisheries Research 96, 102–108 doi.org/10.1016/j.fishres.2008.09.005

Kottelat, M. & Freyhof, J. (2007) Handbook of European freshwater fishes. Kottelat, Cornol, Switzerland and Freyhof, Berlin, Germany.

Kruskal, J. B. (1964) Nonmetric multidimensional scaling: A numerical method. Psychometrika 29, 115–129 doi.org/10.1007/BF02289694

LAVES - Landesamt für Verbraucherschutz und Lebensmittelsicherheit, Dezernat für Binnenfischerei (2011) Vorläufige Rote Liste der Süßwasserfische, Rundmäuler und Krebse in Niedersachsen (Stand 2008). Hannover. [unveröffentlicht]

Lemmens, P., Mergeay, J., de Bie, T., Van Wichelen, J., de Meester, L. & Declerck, S. A. J. (2013) How to Maximally Support Local and Regional Biodiversity in Applied Conservation? Insights from Pond Management. PLoS ONE 8, 1–13 doi.org/10.1371/journal.pone.0072538

Lemmens, P., Declerck, S. A. J., Tuytens, K., Vanderstukken, M. & de Meester, L. (2018) Bottom-Up Effects on Biomass Versus Top-Down Effects on Identity: A Multiple-Lake Fish Community Manipulation Experiment. Ecosystems 21, 166–177 doi.org/10.1007/s10021-017-0144-x

Lyons, J. (1992) The Length of Stream to Sample with a Towed Electrofishing Unit When Fish Species Richness is Estimated. North American Journal of Fisheries Management 12, 198–203 doi.org/10.1577/1548-8675(1992)012<0198:TLOSTS>2.3.CO;2

Magnuson, J. J., Tonn, W. M., Banerjee, A., Toivonen, J., Sanchez, O. & Rask, M. (1998) Isolation vs. extinction in the assembly of fishes in small northern lakes. Ecology 79, 2941–2956 doi.org/10.1890/0012-9658(1998)079[2941:IVEITA]2.0.CO;2

Mantoura, R. F. C. & Llewellyn, C. A. (1983) The rapid determination of algal chlorophyll and carotenoid pigments and their breakdown products in natural waters by reverse-phase high-performance liquid chromatography. Analytica Chimica Acta 151, 297–314 doi.org/10.1016/S0003-2670(00)80092-6

Martin, C. H. & Turner, B. J. (2018) Long-distance dispersal over land by fishes?: extremely rare ecological events become probable over millennial timescales. Proceedings of the Royal Society B: Biological Sciences 25–28 doi.org/10.1098/rspb.2017.2436

Matsuzaki, S. S., Suzuki, K., Kadoya, T., Nakagawa, M. & Takamura, N. (2018) Bottom-up linkages between primary production, zooplankton, and fish in a shallow, hypereutrophic lake. Ecology 0, 1–12 doi.org/10.1002/ecy.2414

De Meester, L., Declerck, S., Stoks, R., Louette, G., Van De Meutter, F., De Bie, T., Michels, E. & Brendonck, L. (2005) Ponds and pools as model systems in conservation biology, ecology and evolutionary biology. Aquatic Conservation: Marine and Freshwater Ecosystems 15, 715–725 doi.org/10.1002/aqc.748

Mehner, T., Diekmann, M., Brämick, U. & Lemcke, R. (2005) Composition of fish communities in German lakes as related to lake morphology, trophic state, shore structure and human-use intensity. Freshwater Biology 50, 70–85 doi.org/10.1111/j.1365-2427.2004.01294.x

Menezes, R. F., Borchsenius, F., Svenning, J. C., Søndergaard, M., Lauridsen, T. L., Landkildehus, F. & Jeppesen, E. (2013) Variation in fish community structure, richness, and diversity in 56 Danish lakes with contrasting depth, size, and trophic state: Does the method matter? Hydrobiologia 710, 47–59 doi.org/10.1007/s10750-012-1025-0

Mollema, P. N. & Antonellini, M. (2016) Water and (bio)chemical cycling in gravel pit lakes: A review and outlook. Earth-Science Reviews 159, 247–270 doi.org/10.1016/j.earscirev.2016.05.006

Molls, F. & Neumann, D. (1994) Fish abundance and fish migration in gravel-pit lakes connected with the river rhine. Water Science & Technology 29, 307–309 doi.org/10.2166/wst.1994.0126

Monk, C. T. & Arlinghaus, R. (2017) Encountering a bait is necessary but insufficient to explain individual variability in vulnerability to angling in two freshwater benthivorous fish in the wild. PLoS ONE 12, 1–25 doi.org/10.1371/journal.pone.0173989 Editor:

Mueller, M., Pander, J., Knott, J. & Geist, J. (2017) Comparison of nine different methods to assess fish communities in lentic flood-plain habitats. Journal of Fish Biology 1–31 doi.org/10.1111/jfb.13333

Oksanen, J., Blanchet, F. G., Friendly, M., Kindt, R., Legendre, P., McGlinn, D., Minchin, P. R., O’Hara, R. B., Simpson, G. L., Solymos, P., et al. (2018) Vegan: Community ecology package: Ordination, Diversity and Dissimilarities. R package version 2.4-6 2018

Olden, J. D., Kennard, M. J., Leprieur, F., Tedesco, P. A., Winemiller, K. O. & García-Berthou, E. (2010) Conservation biogeography of freshwater fishes: Recent progress and future challenges. Diversity and Distributions 16, 496–513 doi.org/10.1111/j.1472-4642.2010.00655.x

Osgood, R. A. (2005) Shoreline Density. Lake and Reservoir Management 21, 125–126 doi.org/10.1080/07438140509354420

Padilla, D. K. & Williams, S. L. (2004) Beyond ballast water:aquarium and ornamental trades as sources of invasive species in aquatic ecosystems. Frontiers in Ecology and the Environment 3, 131–138 doi.org/10.1890/1540-9295(2004)002[0131:BBWAAO]2.0.CO;2

Paller, M. H. (1995) Relationships among number of fish species sampled, reach length surveyed, and sampling effort in South Carolina coastal plain streams. North American Journal of Fisheries Management 15, 110–120 doi.org/10.1577/1548-8675(1995)015<0110:RANOFS>2.3.CO;2

Patoka, J., Bláha, M., Kalous, L. & Kouba, A. (2017) Irresponsible vendors: Non-native, invasive and threatened animals offered for garden pond stocking. Aquatic Conservation: Marine and Freshwater Ecosystems 27, 692–697 doi.org/10.1002/aqc.2719

Persson, L., Diehl, S., Johansson, L., Andersson, G. & Hamrin, S. F. (1991) Shifts in fish communities along the productivity gradient of temperate lakes-patterns and the importance of size-structured interactions. Journal of Fish Biology 38, 281–293 doi.org/10.1111/j.1095-8649.1991.tb03114.x

Pont, D., Crivelli, A. J. & Guillot, F. (1991) The impact of three-spined sicklebacks on the zooplankton of a previuosly fish-free pool. Freshwater Biology 26, 149–163 doi.org/10.1111/j.1365-2427.1991.tb01725.x

R Core Team (2016) R: A language and environment for statistical computing. R Foundation for Statistical Computing, Vienna, Austria. 2016

Radomski, P. J. & Goeman, T. J. (1995) The Homogenizing of Minnesota Lake Fish Assemblages. Fisheries 20, 20–23 doi.org/10.1577/1548-8446(1995)020<0020:THOMLF>2.0.CO;2</0020:THOMLF>

Rahel, F. J. (2000) Homogenization of fish faunas across the United States. Science 288, 854–856 doi.org/10.1126/science.288.5467.854

Rahel, F. J. (2002) Homogenization of Freshwater Faunas. Annual Review of Ecology and Systematics 33, 291–315 doi.org/10.1146/annurev.ecolsys.33.010802.150429

Rahel, F. J. (2007) Biogeographic barriers, connectivity and homogenization of freshwater faunas: It’s a small world after all. Freshwater Biology 52, 696–710 doi.org/10.1111/j.1365-2427.2006.01708.x

Riehl, R. (1991) Können einheimische Fische anhand ihrer Eier durch Wasservögel verbreitet werden? Zeitschrift für Fischkunde 1, 79–83

Riepe, C., Fujitani, M., Cucherousset, J., Pagel, T., Buoro, M., Santoul, F., Lassus, R. & Arlinghaus, R. (2017) What determines the behavioral intention of local-level fisheries managers to alter fish stocking practices in freshwater recreational fisheries of two European countries? Fisheries Research 194, 173–187 doi.org/10.1016/j.fishres.2017.06.001

Ritterbusch, D., Brämick, U. & Mehner, T. (2014) A typology for fish-based assessment of the ecological status of lowland lakes with description of the reference fish communities. Limnologica 49, 18–25 doi.org/10.1016/j.limno.2014.08.001

Santoul, F., Figuerola, J. & Green, A. J. (2004) Importance of gravel pits for the conservation of waterbirds in the Garonne river floodplain (southwest France). Biodiversity and Conservation 13, 1231–1243 doi.org/10.1023/B:BIOC.0000018154.02096.4b

Santoul, F., Gaujard, A., Angélibert, S., Mastrorillo, S. & Céréghino, R. (2009) Gravel pits support waterbird diversity in an urban landscape. Hydrobiologia 634, 107–114 doi.org/10.1007/s10750-009-9886-6

Schälicke, S., Hühn, D. & Arlinghaus, R. (2012) Strukturierende Faktoren der litoralen Fischartengemeinschaften angelfischereilich bewirtschafteter Baggerseen in Niedersachsen. Forschungsbericht des Besatzfisch Projekts, Leibniz-Institut für Gewässerökologie und Binnenfischerei, Berlin, 73 Seiten. (PDF download unter http://www.besatz-fisch.de).

Scheffer, M., Van Geest, G. J., Zimmer, K., Jeppesen, E., Søndergaard, M., Butler, M. G., Hanson, M. A., Declerck, S. & De Meester, L. (2006) Small habitat size and isolation can promote species richness: Second-order effects on biodiversity in shallow lakes and ponds. Oikos 112, 227–231 doi.org/10.1111/j.0030-1299.2006.14145.x

Schurig, H. (1972) Der Baggersee-ein neuer Gewässertyp. Österreichs Fischerei 1, 1–5

Shannon, C. E. (1948) A Mathematical Theory of Communication. Bell System Technical Journal 27, 379–423 doi.org/10.1002/j.1538-7305.1948.tb01338.x

Šmejkal, M., Ricard, D., Prchalová, M., Říha, M., Muška, M., Blabolil, P., Čech, M., Vašek, M., Ju°za, T., Herreras, A. M., et al. (2015) Biomass and abundance biases in European standard gillnet sampling. PLoS ONE 10, 1–15 doi.org/10.1371/journal.pone.0122437

Søndergaard, M., Lauridsen, T. L., Johansson, L. S. & Jeppesen, E. (2018) Gravel pit lakes in Denmark: Chemical and biological state. Science of the Total Environment 612, 9–17 doi.org/10.1016/j.scitotenv.2017.08.163

Staas, S. & Neumann, D. (1994) Reproduction of fish in the Lower River Rhine and connected gravel-pit lakes. Water Science & Technology 29, 311–313 doi.org/10.2166/wst.1994.0127

Strayer, D. L. & Dudgeon, D. (2010) Freshwater biodiversity conservation: recent progress and future challenges. Journal of the North American Benthological Society 29, 344–358 doi.org/10.1899/08-171.1

Strona, G., Galli, P., Montano, S., Seveso, D. & Fattorini, S. (2012) Global-Scale Relationships between Colonization Ability and Range Size in Marine and Freshwater Fish. PLoS ONE 7 doi.org/10.1371/journal.pone.0049465

Theis, S. (2016) Typology of German Angling clubs in relation to means to manage local fisheries. http://www.ifishman.de/publikationen/einzelansicht/188-typology-of-german-angling-clubs-in-relation-to-means-to-manage-to-local-fisheries/

Theis, S., Riepe, C., Fujitani, M. & Arlinghaus, R. (2017) Typisierung von Angelvereinen in Bezug auf den Einsatz von Hegemaßnahmen. AFZ-Fischwaid 3, 18–20

UEPG (2017) European Aggregates Association - Annual Review 2015-2016. Union Européenne des Producteurs de Granulats 1–32 doi.org/10.1016/0038-092X(81)90058-X

Valley, R. D. (2016) Case Study Spatial and temporal variation of aquatic plant abundance: Quantifying change. Journal of Aquatic Plant Management 54, 95–101 doi.org/10.1111/nph.12907

Villéger, S., Blanchet, S., Beauchard, O., Oberdorff, T. & Brosse, S. (2011) Homogenization patterns of the world’s freshwater fish faunas. Proceedings of the National Academy of Sciences 108, 18003–18008 doi.org/10.1073/pnas.1107614108

Völkl, W. (2010) Die Bedeutung und Bewertung von Baggerseen für Fische, Vögel, Amphibien und Libellen: Vereinbarkeit der fischereilichen Nutzung mit den Anforderungen des Naturschutzes. Bezirk Oberfranken - Fachberatung für Fischerei. Bayreuth, 99 Seiten.

Vörösmarty, C. J., McIntyre, P. B., Gessner, M. O., Dudgeon, D., Prusevich, a, Green, P., Glidden, S., Bunn, S. E., Sullivan, C. a, Liermann, C. R., et al. (2010) Global threats to human water security and river biodiversity. Nature 467, 555–561 doi.org/10.1038/nature09549

Whittaker, R. H. (1972) Evolution and Measurement of Species Diversity. Taxon 21, 213–251 doi.org/10.2307/1218190

Wiesner, C., Wolter, C., Rabitsch, W. & Nehring, S. (2010) Gebietsfremde Fische in Deutschland und Österreich und mögliche Auswirkungen des Klimawandels. Bundesamt für Naturschutz-Skripten 279, 192 Seiten.

Winfield, I. J., van Rijn, J. & Valley, R. D. (2015) Hydroacoustic quantification and assessment of spawning grounds of a lake salmonid in a eutrophicated water body. Ecological Informatics 30, 235–240 doi.org/10.1016/j.ecoinf.2015.05.009

Wolter, C. & Röhr, F. (2010) Distribution history of non-native freshwater fish species in Germany: how invasive are they? Journal of Applied Ichthyology 26, 19–27 doi.org/10.1111/j.1439-0426.2010.01505.x

Wright, S. W., Jeffrey, S. W., Mantoura, R. F. C., A, L. C., Bjornland, T., Repeta, D. & Welschmeyer, N. (1991) Improved HPLC method for the analysis of chlorophylls and carotenoids from marine phytoplankton. Marine Ecology Progress Series 77, 183–196 doi.org/10.3354/meps077183

Zhao, T., Grenouillet, G., Pool, T., Tudesque, L. & Cucherousset, J. (2016) Environmental determinants of fish community structure in gravel pit lakes. Ecology of Freshwater Fish 25, 412–421 doi.org/10.1111/eff.12222

Zwirnmann, E., Krüger, A. & Gelbrecht, J. (1999) Analytik im zentralen Chemielabor des IGB. Berichte des IGB. 9, 3–24

